# The spatial signal in area LIP is not an obligatory correlate of perceptual evidence during informed saccadic choices

**DOI:** 10.1101/2021.02.16.431470

**Authors:** Joshua A Seideman, Terrence R Stanford, Emilio Salinas

**Affiliations:** Department of Neurobiology and Anatomy, Wake Forest School of Medicine, 1 Medical Center Blvd., Winston-Salem, NC 27157-1010, USA

**Author notes:** Correspondence should be addressed to: Emilio Salinas, Department of Neurobiology and Anatomy, Wake Forest School of Medicine, 1 Medical Center Blvd., Winston-Salem, NC 27157-1010, USA, www.urgentchoicelab.org. Co-senior authors (Terrence R Stanford, Emilio Salinas). **Disclosures:** The authors declare no conflicts of interest, financial or otherwise. **Author contributions:** JAS, ES and TRS designed the research; JAS collected data; JAS, ES and TRS analyzed data; JAS, ES and TRS wrote the manuscript.

**Keywords:** attention, accumulation of evidence, choice, decision making, eye movement, parietal cortex, perception

## Abstract

The lateral intraparietal area (LIP) contains spatially selective neurons that are partly responsible for determining where to look next and are thought to serve a variety of sensory, motor planning, and cognitive control functions within this role^1,2,3^. Notably, according to numerous studies in monkeys^4,5,6,7,8,9,10,11,12^, area LIP implements a fundamental perceptual process, the gradual accumulation of sensory evidence in favor of one choice (e.g., look left) over another (look right), which manifests as a slowly developing spatial signal during a motion discrimination task. However, according to recent inactivation experiments^13,14^, this signal is unnecessary for accurate task performance. Here we reconcile these contradictory findings. We designed an urgent version of the motion discrimination task in which there is no systematic lag between the perceptual evaluation and the motor action reporting it, and such that the evolution of the subject’s choice can be tracked millisecond by millisecond^15,16,17,18^. We found that while choice accuracy increased steeply with increasing sensory evidence, at the same time, the spatial selection signal in LIP became progressively weaker, as if it hindered performance. In contrast, in a similarly urgent task in which the discriminated stimuli and the choice targets were spatially coincident, this neural signal seemed to facilitate performance. The data suggest that the LIP activity traditionally interpreted as evidence accumulation may correspond to a slow, post-decision shift of spatial attention from one location (where the motion occurs) to another (where the eyes land).

---

The lateral intraparietal area (**LIP**) combines sensory and cognitive information to highlight behaviorally relevant locations or visual features to look at. Although this may involve many sophisticated perceptual operations^3,19,20,21^, the accumulation of sensory evidence (or, more generally, temporal integration) is one of major theoretical importance. First, by some accounts^22,23^, it is an obligatory antecedent to perceptually guided choices regardless of task details, sensory modality, or effector. And second, its manifestation in LIP provides key experimental justification for sequential sampling models, which comprise the most widespread computational framework for reproducing reaction time (**RT**) and accuracy data in deterministic choice tasks^24,25,26,27^. In this framework, the gradual differentiation between spatial locations signaled by LIP corresponds directly to the gradual formation of the perceptual decision^28,29^.

The random-dot motion (**RDM**) discrimination task (Fig. 1a) has been pivotal to this functional interpretation. In it, the subject must look at one of two choice targets to indicate the net direction of motion of a cloud of flickering dots, and in numerous variants of the task^4,5,6,7,8,9,10,11,12^, LIP neurons gradually signal the chosen location while simultaneously reflecting the particulars of the perceptual discrimination. However, in recent inactivation experiments^13,14^, the LIP spatial signal was disrupted with minimal consequence to performance (effects were seen on RT but not on accuracy), consistent with a more indirect relationship between LIP activity and decision formation^29,30^.

**Figure 1.**
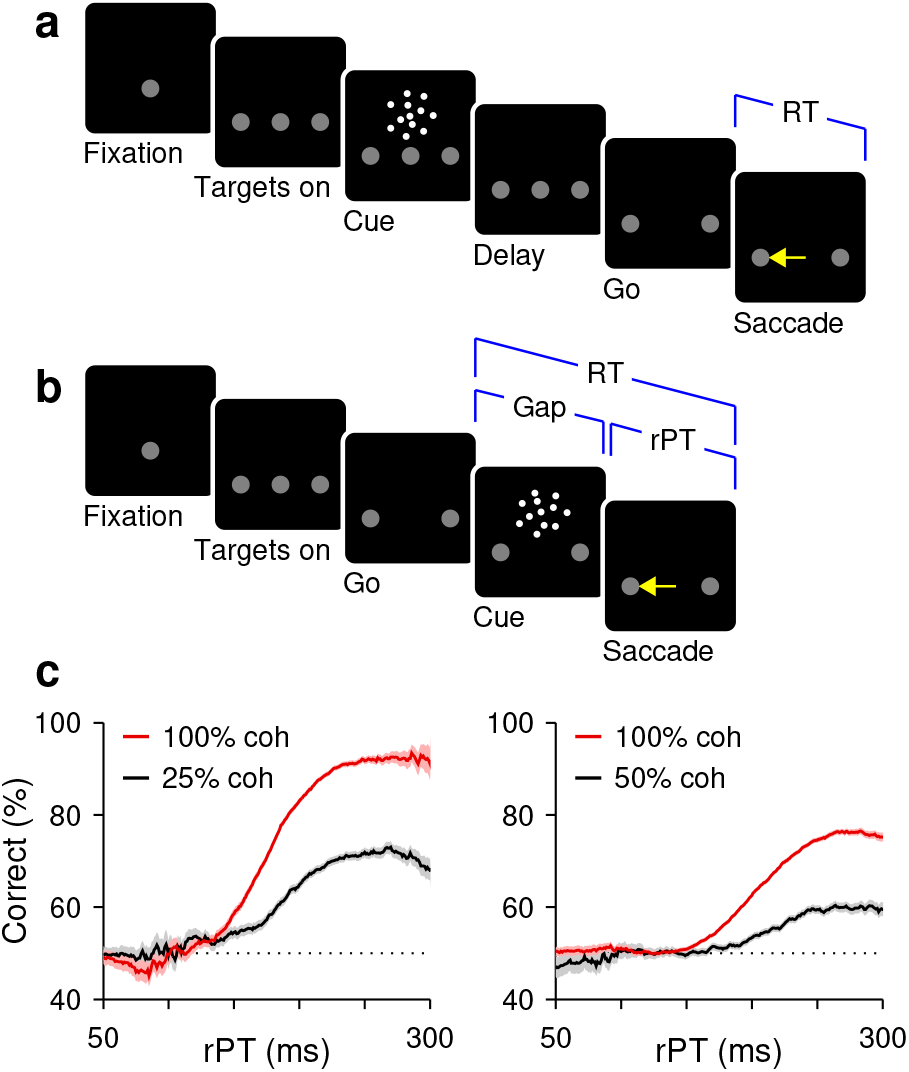
Urgent and non-urgent motion discrimination tasks. Subjects had to report the direction of motion (left or right) of a cloud of flickering dots by looking at one of two peripheral targets. **a**, RDM task (non-urgent). The motion stimulus is presented and evaluated (Cue, 600–1000 ms) well before the go signal (fixation point offset; Go). **b**, CRDM task (urgent). The motion stimulus is presented (Cue) after the go signal (Go), with an unpredictable delay between them (Gap, 0–250) and a limited RT time window for responding (350–425 ms). The perceptual evaluation must occur during the cue-viewing interval (rPT = RT — gap), as the motor plan develops. **c**, Percentage of correct responses as a function of rPT, or tachometric curve. Results are from CRDM behavioral sessions for monkeys C (left) and T (right) for 100% (red; C: 9544, T: 33974 trials) and a lower coherence (black; C: 7909, T: 12066 trials). Shades indicate ± 1 SE from binomial statistics.

We propose a simple yet potentially far-reaching explanation for this puzzling combination of findings: the perceptual evaluation of the motion stimulus occurs more rapidly (∼200 ms) than is generally assumed and may *precede* the LIP differentiation in many instances. So, what appears to be a gradual accumulation of sensory evidence is likely the byproduct of task designs that promote a slow, post-decision shift of attention from one spatial location (where the dots are) to another (where the chosen target is).

This hypothesis makes a stark prediction. Consider the RDM task performed with high urgency, such that perceptual and motor planning processes run concurrently (Fig. 1b, c). This will produce correct trials that are rapid (low RT) but still informed by the motion stimulus. If LIP neurons accumulate evidence, then in those trials they must still differentiate and indicate the impending choice, with stronger evidence yielding stronger differentiation. Alternatively, if the spatial differentiation in LIP occurs after the motion stimulus has been evaluated, its development on such rapid trials will be curtailed, and stronger evidence will not prevent its attenuation or abolition altogether.

## Urgent versus non-urgent choices

To test this prediction, we recorded single-neuron activity in area LIP during two variants of the RDM discrimination task. In the standard, non-urgent version (Fig. 1a), the motion stimulus is presented first (Cue, 600–1000 ms) and is followed by the offset of the fixation point (Go), which means “respond now!” In the urgent or compelled random-dot motion (**CRDM**) discrimination task (Fig. 1b), the order of events is reversed: the go signal is given first, before the stimulus is shown, and the subject must respond within a short time window after the go (350–425 ms). Although the required perceptual judgment is the same, the tasks differ critically in the order in which perceptual and motor processes are engaged. In the former, the saccade can be prepared with relative leisure, after the perceptual evaluation is completed, whereas in the latter, the motor plan is initiated early and the perceptual evaluation must occur while the developing motor plan advances. Under time pressure, saccades can be triggered before, during, or shortly after the perceptual evaluation, and may result in guesses, partially informed, or fully informed choices (Fig. 1c). Perceptual and motor performance are effectively decoupled^15,16,17,18^ (Fig. S1).

Two monkey subjects performed the two choice tasks in interleaved blocks of trials (in addition to single-target tasks traditionally used to characterize LIP activity; Fig. 2a, b). In the standard, non-urgent RDM task, most choices were correct (93% and 84% correct for monkeys C and T at 100% coherence; Fig. S2), and the recorded LIP activity evolved as reported previously^4,5,8,11^ (Fig. 2c). The neurons responded briskly upon presentation of a choice target in the response field (**RF**), continued firing at an elevated rate, and began signaling the choice about 200 ms after the onset of the motion stimulus (Fig. 2d, red arrow), at which point their activity increased for saccades into the RF and decreased for saccades away.

**Figure 2.**
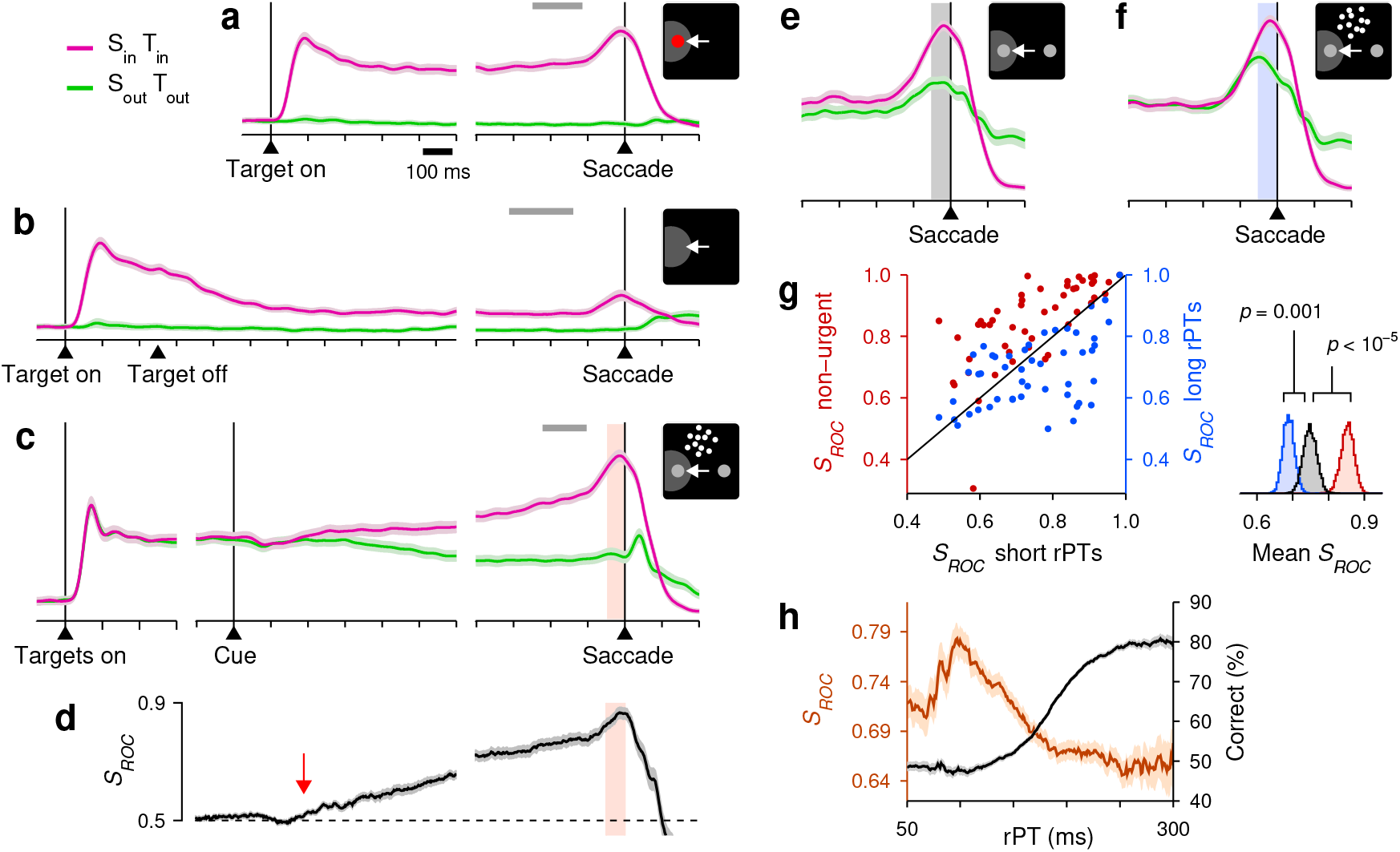
LIP activity in urgent versus non-urgent random-dot motion discrimination. **a**, Responses during visually guided saccades. Traces show normalized firing rate (mean ± 1 SE across cells; *n* = 50) as a function of time for correct trials into (magenta) or away from the cell’s RF (green). Same scales for other panels. The gray bar indicates the go signal range for 90% of the trials. **b**, Responses during memory guided saccades (*n* = 49). **c**, Responses in the non-urgent RDM task (*n* = 51). **d**, Spatial signal magnitude as a function of time for the data in **c** (same time axis). Throughout the article, *S*_*ROC*_ measures the statistical separation between inward and outward responses (Methods). Red arrow marks approximate onset of differentiation. **e, f**, Responses in the CRDM task (*n* = 51) during guesses (**e**, rPT ≤ 150 ms) and fully informed choices (**f**, rPT ≥ 200 ms). **g**, Presaccadic *S*_*ROC*_ for individual neurons (*n* = 51) during guesses (x-axis) and fully informed choices in the CRDM task (right y-axis), and in the non-urgent RDM task (left y-axis). Spike counts for computing *S*_*ROC*_ are from shaded windows in **c**–**f**. Side plot shows bootstrapped distributions of mean values. **h**, Behavioral (black) and neuronal (brown) performance curves from the same CRDM sessions (mean ± 1 SE across trials). *S*_*ROC*_ is from presaccadic spikes pooled across neurons (*n* = 51) and sorted by rPT (bin width = 51 ms). All motion data are for 100% coherence.

To interpret this growing differential signal (quantified by *S*_*ROC*_, Fig. 2d) as an immediate correlate of the perceptual evaluation — one that is causal to the choice — one must assume that the evaluation begins about 200–250 ms after cue onset. And indeed, many experiments are consistent with such a protracted time scale^6,7,8,10,11,24^. However, none of these studies tracked the timecourse of performance explicitly, moment by moment. By doing this, we find that after 250 ms of stimulus viewing time the motion discrimination is essentially over.

### Perceptual and neural discrimination under time pressure

In the CRDM task, the key variable is the amount of time during which the stimulus can be seen and analyzed before movement onset, which we call the raw processing time (**rPT**, computed as RT − gap in each trial; Fig. 1b). Plotting choice accuracy as a function of rPT yields a detailed, high-resolution account of the temporal evolution of the perceptual judgment (Figs. 1c, 2h). According to this ‘tachometric’ curve, in trials with rPT ≲ 140 ms the stimulus is seen so briefly that the motion direction cannot be resolved, which results in uninformed choices, or guesses (∼50% correct). Choice accuracy then rises rapidly after the 150 ms mark, reaching asymptotic performance for rPTs of 200–250 ms. This amount of viewing time is sufficient for evaluating the RDM stimulus and reliably determining its motion direction.

As in other urgent tasks with similar designs^15,16,17,18^, the rPT measured in each trial quantifies the degree to which sensory evidence guided the corresponding choice (or the probability that the choice was guided). Thus, if the differential signal in LIP reflects the amount of evidence accumulated in each trial, then it should be larger for fully informed discriminations (at long rPTs) than for guesses (at short rPTs), and its evolution should parallel the rise of the tachometric curve.

Contrary to this expectation, the recorded LIP activity showed quite the opposite. During performance of the CRDM task, the neural responses favoring each of the two possible eye movements were clearly separated just prior to saccade onset (Fig. 2e, f). Quantitatively, the presaccadic separation was less definitive than that in the non-urgent condition (Fig. 2g, red data), but the urgent differential signal still pointed reliably to the eventual choice. Crucially, however, across the sample of individual neurons recorded in the CRDM task (*n* = 51), the differential signal measured during fully informed, correct choices (rPT ≥ 200 ms) was considerably weaker than that during guesses (rPT ≤ 150 ms; Fig. 2g, blue data, *p* = 0.001, permutation test). More evidence yielded less differentiation. Furthermore, when the presaccadic responses were pooled across neurons and binned by rPT to assess how the spatial signal develops as a continuous function of processing time (Methods), the resulting neurometric curve decreased steadily for rPT *>* 100 ms (Fig. 2h, brown curve) — in sharp contrast to choice accuracy (Fig. 2h, black curve).In the CRDM task, the stronger the influence of perception on the choice, the weaker the observed LIP differentiation.

### Relationship between LIP differentiation and trial outcome

Everything else being equal, the neural encoding of perceptual information upon which choices are made is typically more robust for correct than for incorrect outcomes^31,32,33,34^. This is true across tasks, circuits, and modalities, and should apply to urgent choices too. During short-rPT trials (rPT ≤ 150 ms), the differential response in LIP was identical for correct and incorrect choices (Fig. 3a, gray bars), as anticipated given that those were all guesses. During informed discriminations (rPT *>* 150 ms), however, the differentiation was greater for errors than for correct choices (Fig. 3a, b, blue vs. purple data, *p* = 0.0006, resampling test) — again, opposite to the trend expected from an evidence accumulation process.

**Figure 3.**
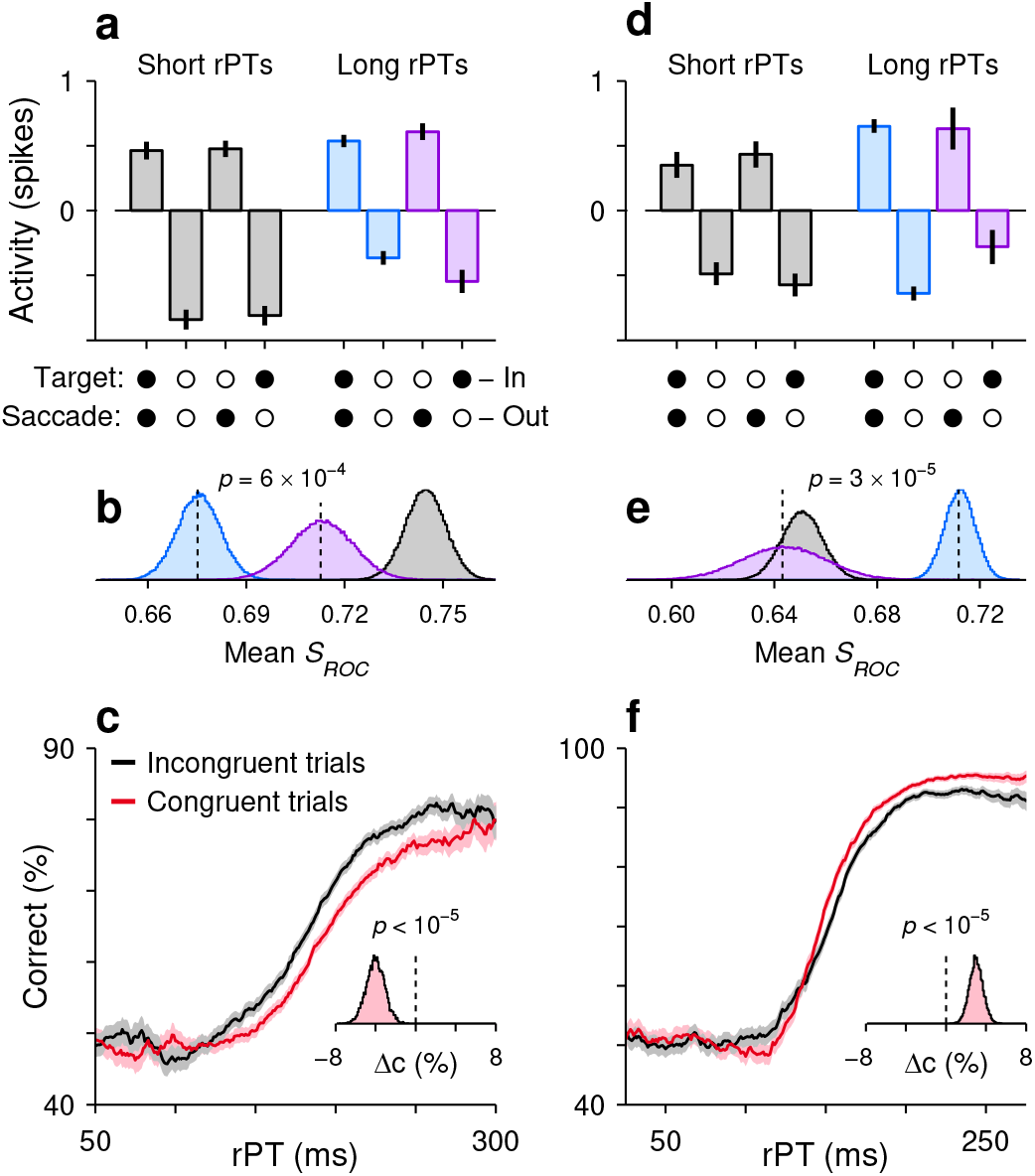
LIP differentiation may help or hinder performance. **a**, LIP activity in the CRDM task during guesses (rPT ≤ 150 ms, gray) and informed choices (rPT *>* 150 ms, blue, purple) sorted by outcome (x-axis). Activity indicates presaccadic spike counts normalized and pooled across neurons (*n* = 51). Data are mean and 95% CIs across trials. **b**, Differential signal magnitudes for the three conditions in **a** indicated by color. Curves are bootstrapped distributions. **c**, Performance in the CRDM task conditioned on neuronal activity. Trials were classified according to their presaccadic spike counts as either congruent (red) or incongruent (black) with strong differentiation. Inset shows bootstrapped distribution for the mean difference in percent correct between curves for rPTs of 130–230 ms. **d**–**f**, As in **a**–**c**, but for the urgent color discrimination task (*n* = 56; in **d, e**, rPT ≤ 125 ms for guesses and rPT *>* 125 ms for informed choices; in **f**, difference evaluated for rPTs between 140–280 ms).

In urgent tasks, the relationship between behavioral performance and single-neuron activity is revealed most effectively by conditioning the former on the latter. First, for a given experimental condition (saccade into or away from the RF), the spike counts collected from a neuron are sorted by magnitude (above vs. below the median), and then performance is compared across the corresponding groups of trials (Fig. S3; Methods). The resulting tachometric curves conditioned on evoked activity reveal if, when, and how the subject’s behavior changes when the recorded neurons fire more or less than average. According to this analysis, performance was consistently worse (*p <* 10^−5^, resampling test) in trials that were congruent with stronger spatial differentiation (Fig. 3c), as if a more robust spatial signal interfered with the urgent motion discrimination.

### Spatial conflict within LIP

Why is the LIP differentiation suppressed in the CRDM task, and more so for informed choices? There are two likely reasons, both brought about by urgency. First, the differential signal is curtailed when it has less time to develop (Fig. 2g, red data), a general effect^18^ consistent with our initial hypothesis. And second, given LIP’s participation in attentional deployment^2,35,36^, the particular geometry of the task must create a spatial conflict: the early motor plan, initiated shortly after the go signal^15,18^, automatically allocates attentional resources to the planned saccade endpoint(s)^37,38,39,40^, but attention should be directed to the RDM stimulus, which defines the perceptually relevant location^13,14^. A spatial competition ensues^35,41^. Evidence of this is plainly manifest in the behavioral CRDM data (Fig. S4).

To investigate the contributions of these two factors, limited time and attentional conflict, we recorded LIP activity from the same monkeys during two versions, urgent and non-urgent, of a discrimination task in which the subject must make an eye movement to the peripheral stimulus that matches the color of the fixation point^42^ (Fig. S5). The key difference here is that the conflict described above is eliminated: the relevant color cues are found at the choice targets, and deploying attention/perceptual resources to them should be of benefit, if not a necessity, to the required discrimination (Fig. S6).

During the non-urgent color-matching task, the sampled neurons (which again exhibited characteristic LIP response features; Fig. 4a, b) differentiated saccades into versus away from the RF (Fig. 4c, d) slightly earlier than during the standard RDM task (Figs. 2d, 4d, arrows). But, overall, under relaxed, non-urgent conditions, the evoked spatial signal developed with comparable timecourse and strength in the motion- and color-based tasks, in spite of their distinct spatial and feature requirements. Under time pressure, though, the comparison across tasks was striking. During the urgent color-matching task, the differential response in LIP was larger for informed than uninformed discriminations (Fig. 4e–g); its magnitude increased over time in parallel with the monkeys’ choice accuracy (Fig. 4h); it was weaker for errors than correct choices during informed trials (Fig. 3d, e); and it acted as if to improve the monkeys’ performance (Fig. 3f). In this case, the greater the influence of perception on the choice, the stronger the spatial signal observed in LIP.

**Figure 4.**
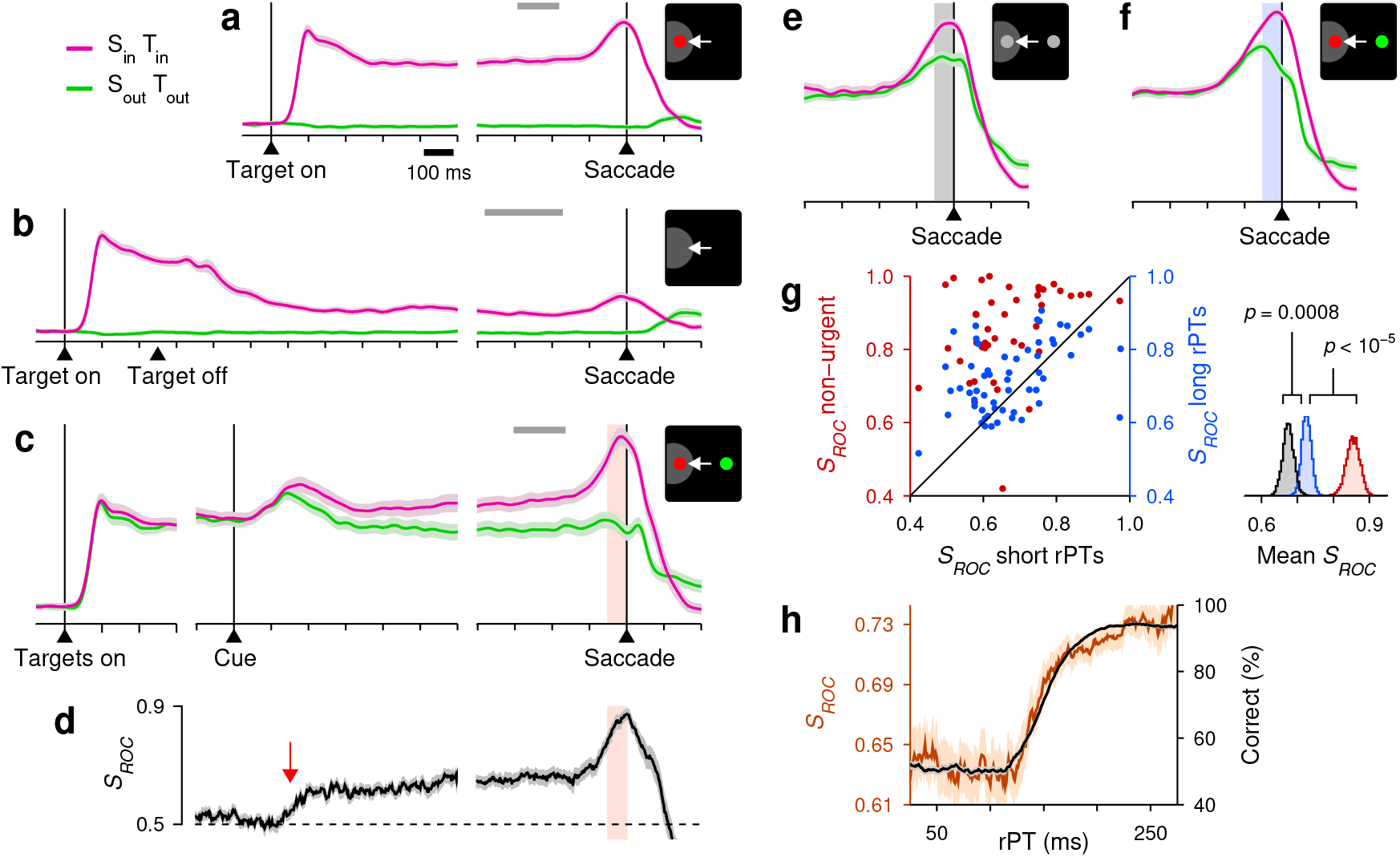
LIP activity in urgent versus non-urgent color discrimination. Same format and conventions as in Fig. 2. Data are from *n* = 56 sampled neurons, except in **c, d**, and **g** (red data), for which *n* = 43. In **e**, rPT ≤ 125 ms. In **f**, rPT ≥ 175 ms.

These results in the color-matching task experiment confirm that an informed spatial signal can emerge very rapidly in LIP^43,44^. They show that time pressure alone does not necessarily abolish or reverse the expected correlation between evidence and LIP differentiation, and so cannot explain the CRDM results. Rather, the data suggest that the anticorrelation between CRDM performance and LIP spatial signal strength results from urgency exacerbating a spatial conflict between the perceptually relevant location and the saccade endpoint. In this case, early selection of the saccade target corresponds to attention being diverted away from the location of the dots during a brief but critical period of time when the motion stimulus is being evaluated.

Notably, an early bias favoring choices into the RF is visible in the CRDM data (Fig. 2e), but this simply reflects a consistent preference for the initial guess that is required of the subjects in every urgent trial. Such consistency is of little consequence to the perceptual evaluation^9,15^. Indeed, the results did not change qualitatively when this bias was eliminated on a trial-by-trial basis (Fig. S7), nor when it was either enhanced or suppressed by suitable selection of experimental sessions (Figs. S8) or recorded trials (Fig. S9). Also, for both the motion- and color-based tasks the results were robust with respect to the subjects’ performance level (Fig. S10), the criteria used for including/excluding neurons (Figs. S11, S12, S13), and how the effects were quantified (Fig. S14).

## Conclusions

The highly robust target selection seen during non-urgent conditions (RDM task) would lead one to conclude, as have countless past studies, that LIP differentiation is an obligatory, causal antecedent to perceptually informed choices, and that greater differentiation implies more or stronger perceptual evidence. Yet, for equally informed choices made urgently (CRDM task), the spatial signal was markedly diminished, and it decreased with increasing evidence. While counterintuitive, this result remains consistent with a representation of spatial priority^2,35,41^.

In general, the differential activation of oculomotor neurons denotes the potential for selection of different relevant locations besides the eventual saccade target. These may contain reward information, a visual cue, a symbolic instruction, or a salient distracter^2,35,36,45,46,47^. Such versatility is the hallmark of attention-related activity. During motion discrimination, the location that is relevant to the perceptual evaluation is where the dots are, and indeed, inactivation experiments indicate that the LIP neurons with RFs covering the dots are precisely the ones that matter the most for performance^13,14^. Under urgent conditions (CRDM task), stronger differentiation between the two perceptually irrelevant choice-target locations would denote a firmer attentional commitment, and likely a stronger conflict with the RFs covering the dots. However, when the urgency requirement is relaxed (standard RDM task), attention can be deployed to the location of the dots even before motion onset, and can remain there as long as necessary to then shift to the chosen saccade target. The focus on the dots need not be long (≲ 100 ms, considering the time to transition from chance to asymptotic performance; Figs. 1c, 2h), and the timecourse and magnitude of the shift may still depend on the strength of the sensory evidence. If so, the resulting post-perceptual differentiation may appear causal to the choice.

## Methods

### Subjects and setup

All experimental procedures were conducted in accordance with NIH guidelines, USDA regulations, and the policies set forth by the Institutional Animal Care and Use Committee (IACUC) of Wake Forest School of Medicine. The subjects in this experiment were two adult male rhesus monkeys (Macaca mulatta) weighing between 8.5 and 11 kg. For each animal, an MRI-compatible post (Crist Instruments, MD, USA) was implanted on the skull while under general anesthesia. The post served to fix the position of the head during all experimental sessions. Following head-post implantation, both subjects were trained to perform oculomotor response tasks in exchange for water reward. After reaching a criterion level (*>* 75% accuracy for each task), craniotomies were made and recording cylinders (Crist Instruments, MD, USA) were placed over the left LIP of each monkey (monkey C: left hemisphere; monkey T: left and right hemispheres; stereotactic coordinates: 5 posterior, 12 lateral^48,49^) while under general anesthesia. Neural recordings commenced after a 1-2 week recovery period following cylinder placement.

### Behavioral and neurophysiological recording systems

Eye position was monitored using an EyeLink 1000 Plus infrared tracking system (SR Research; Ottawa, Canada) at a sampling rate of 500 or 1000 Hz. For sessions in which dot-motion tasks were performed, all gaze-contingent stimulus presentation and reward delivery were controlled using Psychtoolbox^50,51^ version 2.0; for all other sessions, gaze-contingent stimulus presentation and reward delivery were controlled via a custom-designed PC-based software package (Ryklin Software). Visual stimuli were presented on a Viewpixx/3D display (Vpixx Technologies, Quebec, Canada; 1920 × 1080 screen resolution, 120 Hz refresh rate, 12 bit color) placed 57 cm away from the subject. Viewing was binocular. Neural activity was recorded using single tungsten microelectrodes (FHC, Bowdoin, ME; 2–4 MΩ impedance at 1 kHz) driven by a hydraulic microdrive (FHC). A Cereplex M headstage (Blackrock Microsystems, Utah, USA) filtered (0.03 Hz – 7.5 kHz), amplified, and digitized electrical signals, which were then sent to a Cereplex Direct (Blackrock Microsystems) data acquisition system. Single neurons were isolated online based on amplitude criteria and/or waveform characteristics.

### Behavioral tasks

Three design elements are the same for all the tasks. (1) Each trial begins with presentation of a central spot and the monkey fixating it for 300–800 ms. (2) The offset of the fixation spot is the go signal that instructs the monkey to make a saccade. (3) To yield a reward (drop of liquid), the saccade must be to the correct location and must be initiated within an allotted RT window. The RT is always measured as the time elapsed between fixation offset and saccade onset (equal to the time point following the go signal at which the eye velocity first exceeds a criterion of 25 deg/s). In non-urgent tasks the monkey is allowed to initiate an eye movement within 600 ms of the go signal, whereas in urgent tasks this must happen within 350–425 ms.

### Visually- and memory-guided saccade tasks

Two standard single-target tasks were used to characterize the visuomotor properties of LIP neurons. In both tasks, after the monkey fixates, a peripheral target is presented (Target on) either within or diametrically opposed to the RF of the recorded neuron. For the delayed visually-guided saccade task, after a variable delay (500–1000 ms), the fixation spot disappears (Go) and the monkey is required to make a saccade to the peripheral target. For the memory-guided saccade task, after being displayed for 250 ms, the peripheral target is extinguished (Target off) and the monkey is required to maintain fixation throughout a subsequent delay interval (500–1000 ms). After this memory interval, the fixation spot disappears (Go) and the monkey is required to make a saccade to the remembered target location.

### Non-urgent RDM motion discrimination task

This two-alternative task (Fig. 1a) is similar to previous implementations of the RDM discrimination task^4,5,8,11^. Upon fixation and after a short delay (300–500 ms), two gray stimuli, the potential targets, are presented (Targets on), one in the RF and one diametrically opposed. After a delay (250–750 ms), a cloud of randomly moving dots appears in the center of the screen or just above the fixation point for 600–1000 ms (Cue). Then, after another delay period (300–500 ms; Delay), the fixation spot is extinguished (Go), which instructs the monkey to make a choice. If the saccade is to the stimulus in the direction of the dot motion (and is made within 600 ms), the monkey obtains a liquid reward. The direction of motion, toward one choice target or the other, is assigned randomly from trial to trial. The difficulty of the task varies with stimulus coherence, which is the percentage of dots that move in a consistent direction across video frames. Monkeys worked with coherence values of 100%, 50%, 25%, 6% and 3%, but the neural data were recorded at 100% (Fig. S2).

### Compelled random-dot motion discrimination task

The CRDM task (Fig. 1b) is an urgent version of the RDM discrimination task just described. The geometry, reward size, and stimuli are the same; only the temporal requirements are different. In this case, the monkey fixates, the two peripheral gray stimuli are shown (Targets on), and after a delay (250–750 ms), the go signal is given (Go), urging the subject to respond as quickly as possible (within 350–425 ms). At this point in the trial, however, no information is available yet to guide the choice.That information, conveyed by the cloud of flickering dots, is revealed later (Cue), after an unpredictable amount of time following the go (Gap; 0–250 ms). Subjects are tasked with looking to the peripheral choice alternative that is congruent with the net direction of motion of the dots (Saccade).

On each trial, the raw processing time, or rPT, is the maximum amount of time that is potentially available for seeing and evaluating the motion stimulus. It is the time interval between cue onset and saccade onset (rPT = RT − gap). We refer to it as ‘raw’ because it includes any afferent or efferent delays in the circuitry^15^. Gap values (0–250 ms) varied randomly from trial to trial and were chosen to yield rPTs covering the full range between guesses and informed choices.

### Non-urgent color discrimination task

In this task (Fig. S5a), the color of the central fixation spot (red or green) defines the identity of the eventual target. Upon fixation and after a short delay (300–800 ms), two gray stimuli, the potential targets, are presented (Targets on), one in the RF and one diametrically opposed. After a delay (250–750 ms), one of the gray stimuli changes to red and the other to green (Cue). After a cue viewing period (500–1000 ms), the fixation spot is extinguished (Go), which instructs the monkey to make a choice. If the ensuing saccade is to the stimulus that matches the color of the prior fixation spot and is made within 600 ms, the monkey obtains a reward. Colors and locations for target and distracter are randomly assigned in each trial.

### Urgent color discrimination task

This task (Fig. S5b), also referred to as the compelled-saccade task^15,16,18,42^, requires the same red-green discrimination as in the easier non-urgent version. In this case, after the monkey fixates (300–800 ms) and the two gray stimuli in the periphery are displayed (Targets on; 250–750 ms), the fixation spot disappears (Go). This instructs the monkey to make a choice, although the visual cue that informs the choice (one gray spot turning red and the other green; Cue) is revealed later, after an unpredictable period of time following the go signal (Gap; 0–250 ms). To obtain a reward, the monkey must look to the peripheral stimulus that matches the color of the initial fixation spot (Saccade) within the allowed RT window (350–425 ms). As with the CRDM task, the key variable that determines performance is the rPT.

### Tachometric curves and rPT intervals

All data analyses were performed in Matlab (The MathWorks, Natick MA). To compute the tachometric curve and rPT distributions, trials were grouped into rPT bins of 51 ms, with bins shifting every 1 ms. Numbers of correct and incorrect trials were then counted within each bin.

When parsing trials into short and long rPT time bins (Figs. 2e–g, 3a, b, d, e, 4e–g), we considered the distributions of processing times from all the recording sessions in each task. The threshold for guesses (rPT ≤ 150 for the CRDM task; rPT ≤ 125 ms for the color task) corresponded to the point at which the fractions of correct and incorrect trials started diverging steadily with rPT. Trials above this cutoff were considered informed, and trials above this cutoff plus 50 ms, which brought the fraction correct about 75% of the way from chance to asymptotic, were considered fully informed. The results depended minimally on the exact cutoffs used.

Tachometric curves conditioned on neuronal activity (Fig. 3c, f) were computed as follows. First, for each neuron, spike counts from a presaccadic window (−50:0 ms, aligned on saccade) were collected and sorted into two conditions, saccade-in (*S*_*in*_) and saccade-out (*S*_*out*_) choices. The trials in each condition were then split into two groups, with spike counts below the median for the condition, or with spike counts at or above it. Four groups of trials resulted: *S*_*in*_ high firing, *S*_*in*_ low firing, *S*_*out*_ high firing, and *S*_*out*_ low firing. Data from all the neurons in a sample were aggregated, and a tachometric curve was generated for each group (Fig. S3). The first and last groups are congruent with a strong spatial signal, whereas the other two are incongruent. Because the results were consistent for *S*_*in*_ and *S*_*out*_ conditions (Fig. S3), congruent and incongruent trials were combined across conditions.

For the CRDM data, differences between tachometric curves conditioned on low versus high firing were quantified and evaluated for significance (see below) for rPTs of 130–230 ms. This same range was used for all such analyses, regardless of how the data were parsed. For the urgent color discrimination data, the corresponding range was 140–280 ms.

### Characterization of neural activity

On line, RF location was determined from activity levels measured around the time of saccade onset during performance of the visually- or memory-guided saccade task. All neurons included in the current study (*n* =51 for CRDM task, *n* =56 for urgent color discrimination task) were significantly activated during performance of the urgent tasks, both in response to visual stimuli pre-sented in their RF (window: 20:150 ms, aligned on targets on) as well as prior to saccades executed into their RF (window: −100:0 ms, aligned on saccade) relative to respective baseline measures. In addition, all neurons included exhibited significant delay period activity in the visually- and/or memory-guided saccade tasks. For all such determinations, significance (*p <* 0.01) was calculated numerically via permutation tests^52^ in which the two group labels (e.g., ‘baseline’ and ‘response period’) were randomly permuted. These physiological response properties (i.e., visual, delay period, and presaccadic activation) are characteristic of LIP neurons that project directly to saccade production centers^53^ (i.e., the superior colliculus).

Some additional neurons that were recorded and fully characterized (15 in the CRDM experiment, 26 in the color-based) were excluded from the studied samples for any of the following reasons: they had no significant visual or memory activity in the single-target tasks; they were not significantly activated presaccadically; or their spatial preference for contralateral/ipsilateral stimuli either was ambiguous or clearly flipped between different tasks. Importantly, though, except for small quantitative variations, all results were essentially identical with inclusion of all such neurons (Fig. S11).

For each neuron, continuous firing rate traces, or spike density functions, were generated by aligning the recorded spike trains to relevant task events (e.g., cue onset, saccade onset), convolving them with a gaussian kernel (*σ* =15 ms), and averaging across trials. Normalized population traces (as in panels a–c, e, f of Figs. 2, 4) were generated by dividing each cell’s response curve by its maximum firing rate value and then averaging across cells. For each cell, this maximum rate was calculated from the recorded urgent trials (motion- or color-based) and was used to normalize the population traces for all other tasks.

### ROC analyses and neurometric curves

The magnitude of spatial differentiation, or *S*_*ROC*_, was used to quantify the degree to which LIP neurons were differentially activated in *S*_*in*_ versus *S*_*out*_ choices. This measure corresponds to the accuracy with which an ideal observer can classify data samples from two distributions (of responses in *S*_*in*_ and *S*_*out*_ trials, in this case), and is equivalent to the area under the receiver operating characteristic, or ROC, curve^54,55^. Values of 0.5 correspond to distributions that are indistinguishable (chance performance, full overlap), whereas values of 0 or 1 correspond to fully distinguishable distributions (perfect performance, no overlap). Here, *S*_*ROC*_ > 0.5 always indicates higher activity for saccades into the RF than away from the RF. Presaccadic *S*_*ROC*_ values (Figs. 2g, h, 3b, e, 4g, h) were computed using spike counts measured prior to choice onset (−50:0 ms, relative to saccade onset) and sorted according to trial outcome.

For the urgent tasks, continuous neurometric functions comparable to the behavioral tachometric curves (Figs. 2h, 4h) were generated by first pooling the data across neurons and then calculating *S*_*ROC*_ as a function of rPT (bin width = 51 ms, shifted every 1 ms). The pooling involved two steps. First, the presaccadic spike counts of each neuron were normalized by subtracting a constant, *θ*, that was cell-specific, and then the normalized spike counts from all the neurons were sorted into two groups, for *S*_*in*_ and *S*_*out*_ trials. The pooled *S*_*ROC*_ compared responses from these two pooled distributions within each rPT bin (see Fig. S15 for an example). For each neuron, the constant *θ* was equal to (*m*_*in*_ + *m*_*out*_)*/*2, where *m*_*in*_ and *m*_*out*_ are the mean spike counts for *S*_*in*_ and *S*_*out*_ trials. Other normalization schemes produced qualitatively similar trends. This procedure, pooling the data first and then computing *S*_*ROC*_, generated more precise results than the reverse, i.e., first computing *S*_*ROC*_ for each cell and then averaging across cells. However, the latter alternative produced qualitatively consistent results (Fig. S14). We stress that, although the pooled *S*_*ROC*_ values that make up the neurometric curve vary with rPT, they were always based on spike counts measured just prior to the saccade.

For the non-urgent tasks (Figs. 2d, 4d), continuous *S*_*ROC*_ values were again computed by dividing time into sliding bins (bin width = 50 ms, shifted every 1 ms). For each bin, the spikes counted for each neuron in each condition (*S*_*in*_ and *S*_*out*_ trials) were used to calculate that cell’s *S*_*ROC*_, and then values were averaged across cells. Pooling was unnecessary in this case because more data were available in each time bin, but the results with pooling were very similar.

### Statistical tests

Effect sizes for mean *S*_*ROC*_ values were computed by bootstrapping^56,57^; that is, by repeatedly resampling the underlying data with replacement (10^4^ – 10^5^ iterations) and recomputing the mean *S*_*ROC*_ each time. In Figs. 2g, 4g (insets), the resampling was over neurons; in Fig. 3b, e, it was over trials in the two pooled distributions (for *S*_*in*_ and *S*_*out*_ conditions). Effect sizes for other quantities (e.g., Δ*c* in Fig. 3c, f) were also calculated through bootstrapping. Having generated these effect-size distributions for any two conditions (e.g., correct vs. incorrect choices, or long vs. short rPTs), we could calculate from them a significance value for the mean difference. Instead, however, for any relevant comparison between two conditions, the p-value of the difference was calculated separately using a permutation test^52^ for paired data or an equivalent resampling test for non-paired data, as these tests provide slightly more accurate and specific comparisons against the null hypothesis (of no difference between the distributions from which the two data sets originated). For example, to compare the mean *S*_*ROC*_ for short-versus long-rPT trials (Figs. 2g, 4g, insets), we randomly permuted the ‘short’ and ‘long’ labels for each neuron and recomputed the difference between *S*_*ROC*_ means 10^5^ times. All reported significance values were calculated similarly, via permutation or resampling tests (one-sided).

## Abbreviations

CRDM: compelled random dot motion
RDM: random dot motion
RF: response field
rPT: raw processing time
RT: reaction time
SD: standard deviation
SE: standard error

---

**Figure S1.**
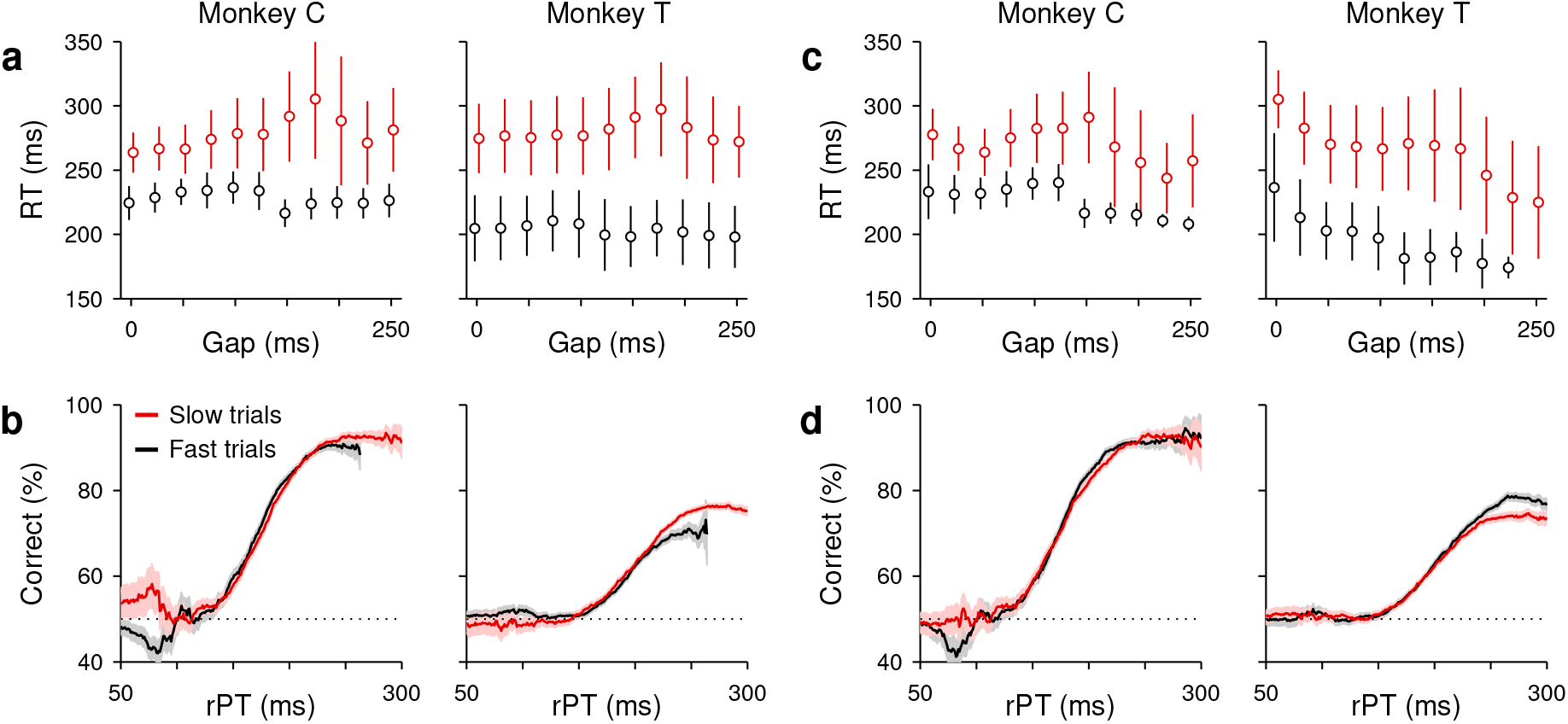
Perceptual and motor performance are decoupled in the CRDM task. For each monkey, trials at 100% coherence were sorted into two groups, slow (red data) and fast (black data). These groups were defined in two ways. In Method 1, slow and fast trials were simply those with RTs above and below the overall median RT, respectively. In Method 2, trials were first sorted into non-overlapping rPT bins (20 ms width), and then the trials in each bin were split into slow and fast according to the median RT of that bin. **a, b**, Mean RT ± 1 SD as a function of gap (**a**) and percentage of correct choices ± 1 SE as a function of processing time, or tachometric curve (**b**), for the slow and fast trials obtained with Method 1. **c, d**, As in **a, b**, but for the slow and fast trials obtained with Method 2. All results are from the CRDM behavioral sessions; same 100% coherence data as in Fig. 1c. In spite of the large differences in RT, the fast and slow trials yielded tachometric curves that were largely indistinguishable. This shows that, under urgent conditions, perceptual performance (response accuracy) during motion discrimination can be reliably quantified independently of motor performance (response speed).

**Figure S2.**
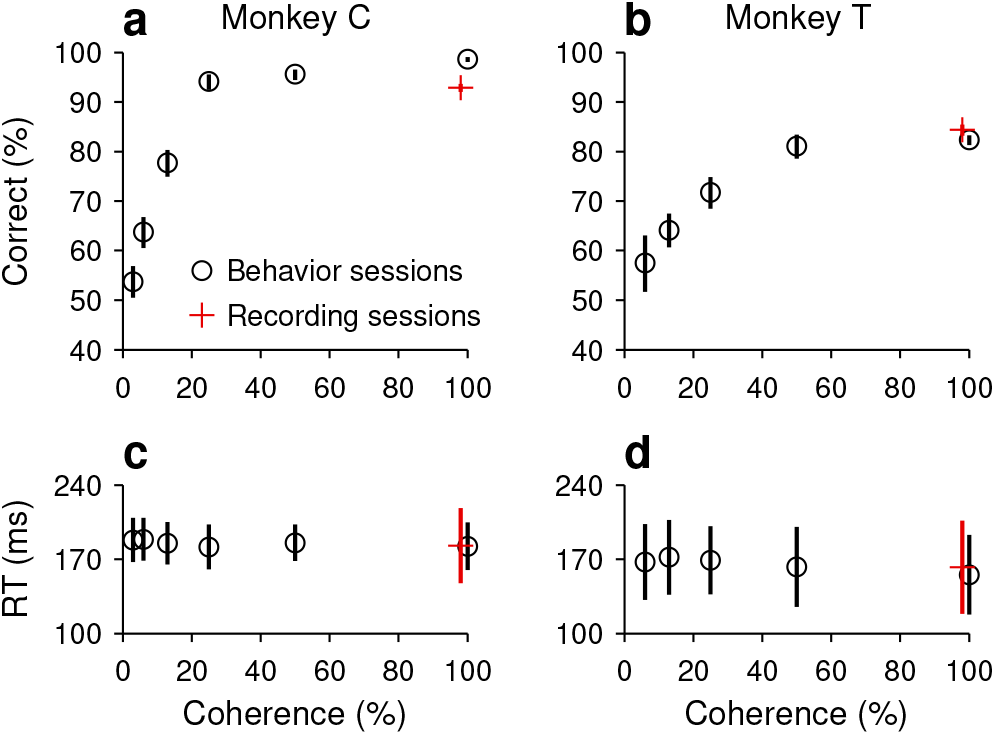
Performance in the non-urgent RDM task. **a**, Percentange of correct choices as a function of stimulus coherence. Data are from monkey C collected during behavioral sessions (black circles, *n* = 7363 trials) or during the recording sessions (red cross, *n* = 8685 trials). Error bars indicate 95% confidence intervals. **b**, Mean RT ±1 SD across trials. **c, d**, As in **a, b**, but for monkey T (black circles, *n* = 4547 trials; red cross, *n* = 3952 trials).

**Figure S3.**
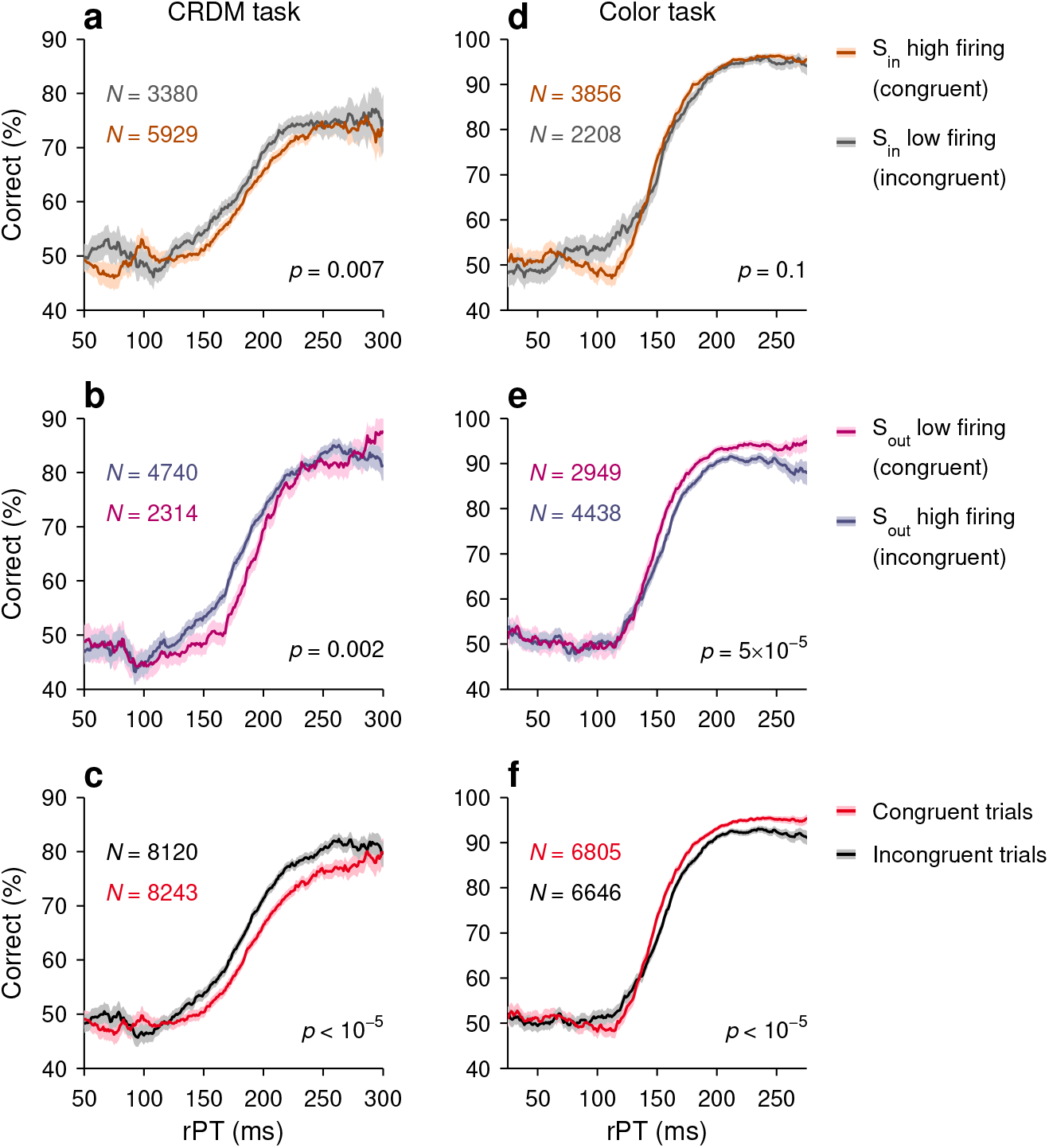
Behavioral performance in the urgent tasks conditioned on LIP neuronal activity. **a**, Tachometric curves from all the CRDM trials in which the outcome was a saccade into the recorded cell’s RF (*S*_*in*_). For each neuron (*n* = 51), *S*_*in*_ trials were split according to whether the response was at or above the median (brown curve, high firing), or below the median (gray curve, low firing). The response was the spike count elicited in the 50 ms immediately preceding the onset of the saccade. High firing in *S*_*in*_ trials is congruent with a strong spatial signal, whereas low firing is incongruent. **b**, Tachometric curves from all the CRDM trials in which the outcome was a saccade away from the recorded cell’s RF (*S*_*out*_). For each neuron, *S*_*out*_ trials were split according to whether the response was at or above the median (blue curve, high firing), or below the median (purple curve, low firing). Low firing in *S*_*out*_ trials is congruent with a strong spatial signal, whereas high firing is incongruent. **c**, Results combining all congruent and incongruent trials across conditions. **d**–**f**, As in **a**–**c**, but for the urgent color discrimination task (*n* = 56). Significance values shown are from resampling tests on the mean difference between each pair of curves (Methods). This difference was evaluated for rPTs of 130–230 ms for the CRDM data and for rPTs of 140–280 ms for the urgent color discrimination data (these ranges apply to all comparisons between conditioned curves in this and other figures). *N* indicates number of trials. Data in **c, f** are the same as in Fig. 3c, f. Note that the results are consistent between *S*_*in*_ and *S*_*out*_ conditions.

**Figure S4.**
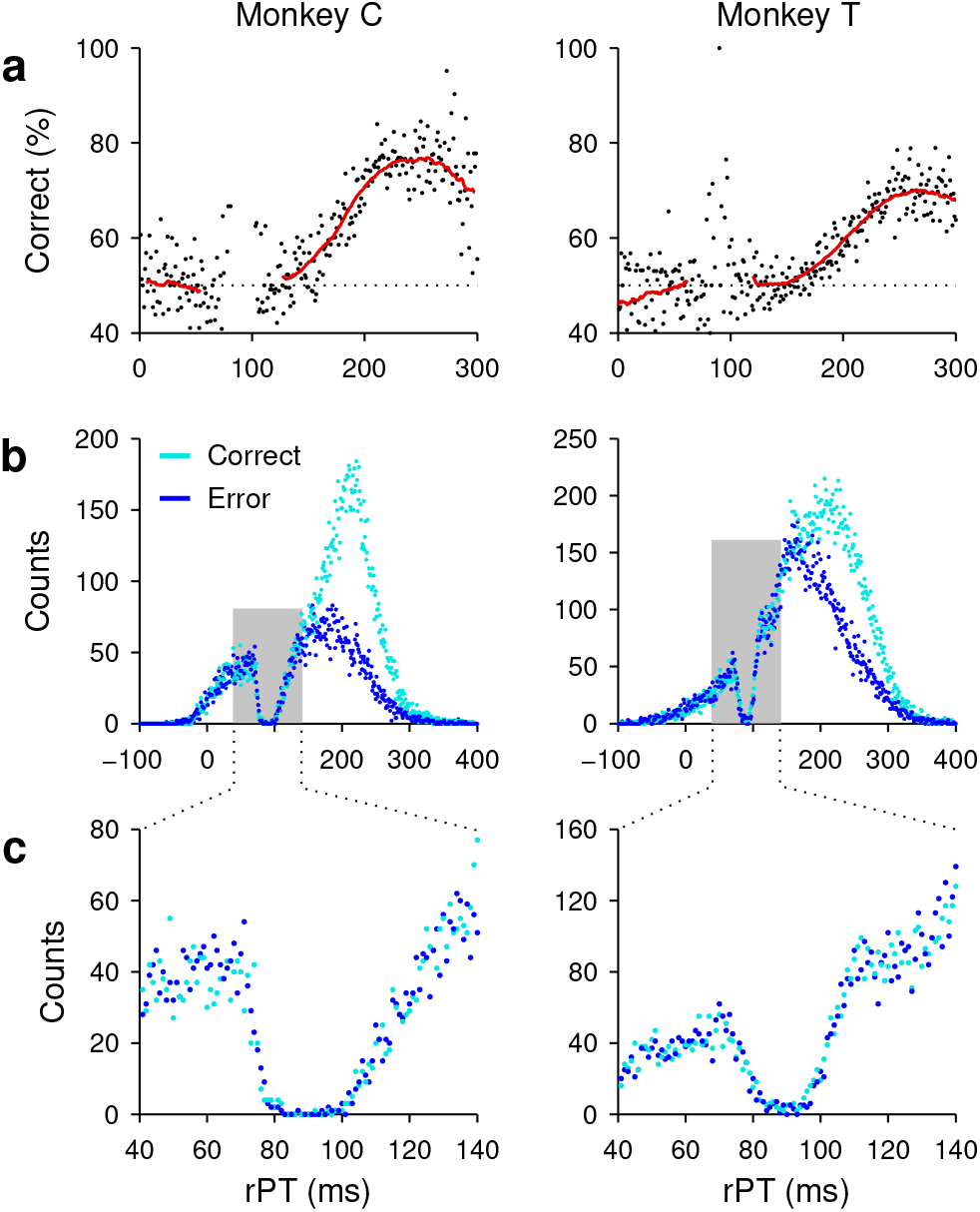
Evidence of spatial conflict in the CRDM task. **a**, Percentange of correct choices as a function of raw processing time. The data comprise all trials from the behavioral sessions of monkey C (left column) and monkey T (right column), and include all coherences. Trials (*n* = 30473 for monkey C, *n* = 52945 for monkey T) were sorted into 50 ms bins (red curve, bins shifted every 1 ms), as done for tachometric curves shown in other figures, or into 1 ms bins (black dots). **b**, Processing time distributions for correct (cyan) and incorrect (blue) choices from the same trials in **a**, sorted into 1 ms bins. **c**, Enlarged view of the data in **b** between 40 and 140 ms of rPT. Note the prominent dip in the number of events at 80–100 ms. We interpret the lack of saccades during this narrow interval as an interruption of the ongoing motor plans due to the onset of the moving-dot stimulus. This is entirely consistent both with the capture of exogenous attention by a salient stimulus^17,35,101^ and with the related phenomenology of saccadic inhibition^102,103,104^. The timing of this dip (∼90 ms after cue onset) is also consistent with a slight decrease in LIP activity often observed^4,5,8,11^ in the non-urgent RDM task (Fig. S6a). For a brief moment, the cue-driven activation at the location of the random dots is in intense conflict with the oculomotor activity that generates saccades to the choice targets.

**Figure S5.**
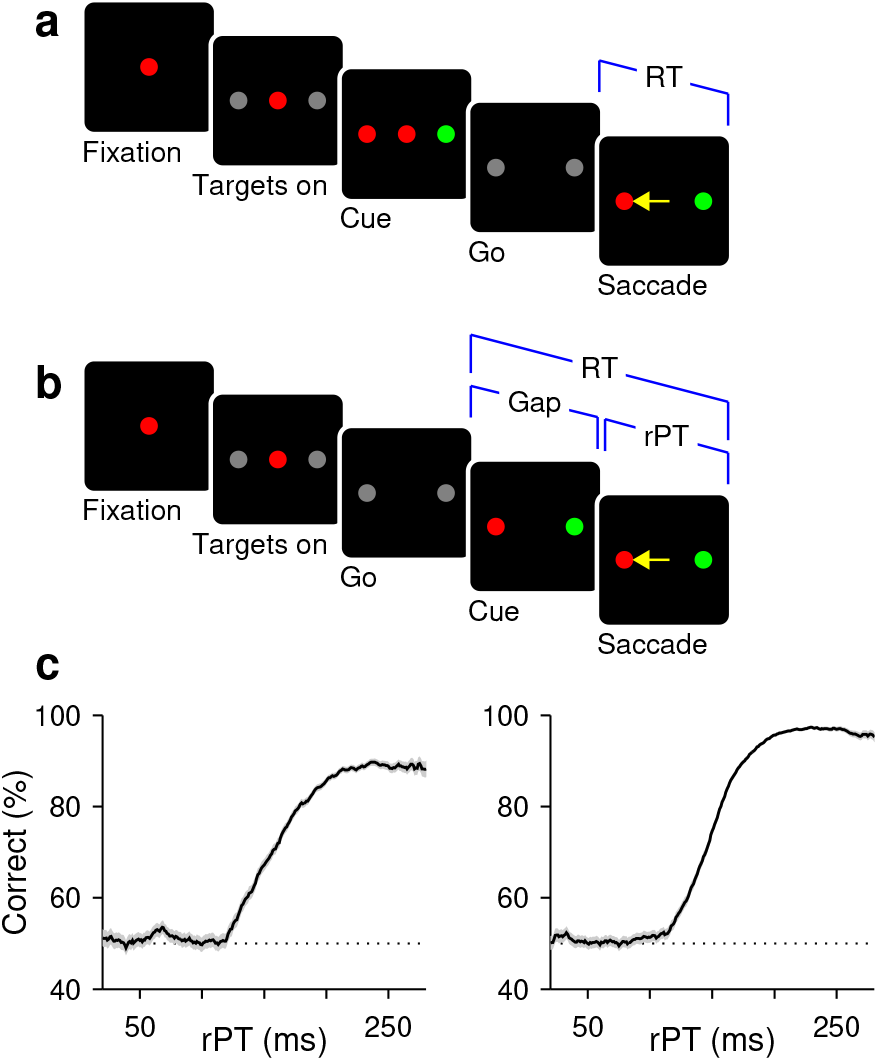
Urgent and non-urgent color-discrimination tasks. Subjects had to look to the peripheral target that matched the color of the fixation point. **a**, Easy, non-urgent task. The color stimuli are presented (Cue, 500–1000 ms) and evaluated well before the go signal (fixation point offset; Go). **b**, Urgent task. The color stimuli are presented (Cue) after the go signal (Go), with an unpredictable delay between them (Gap, 0–250) and a limited RT time window for responding (350–425 ms). The perceptual evaluation must occur during the cue-viewing interval (rPT = RT − gap), as the motor plan develops. **c**, Percentage of correct responses as a function of rPT, or tachometric curve. Results are from all the recording sessions during which the urgent color-discrimination task was performed by monkeys C (left, *n* = 7330 trials) and T (right, *n* = 10745 trials). Shades indicate ± 1 SE from binomial statistics.

**Figure S6.**
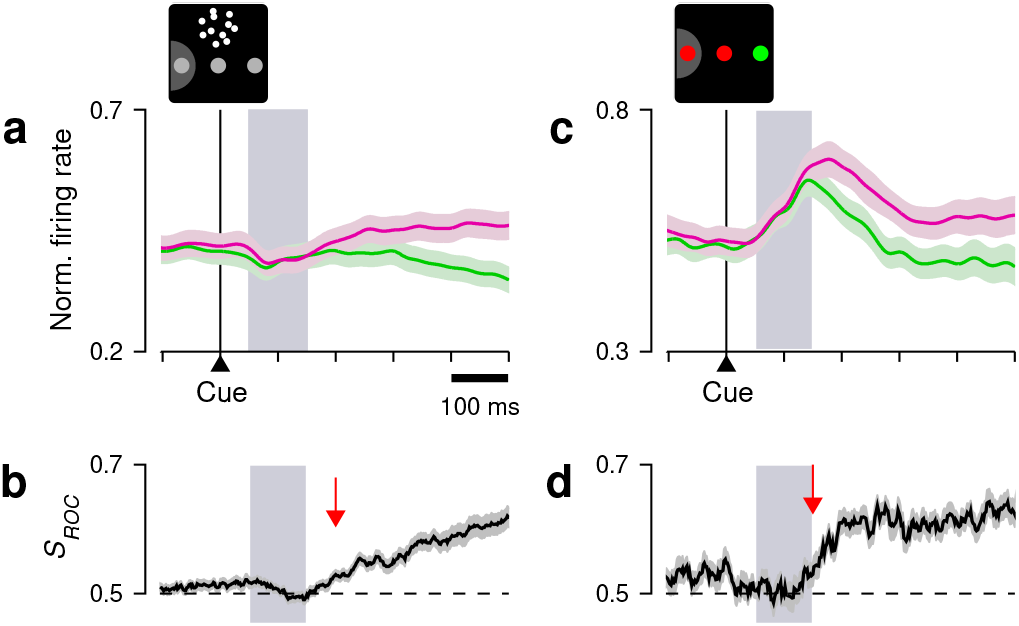
Cue-driven LIP responses in the non-urgent motion- and color-based discrimination tasks are indicative of the deployment of spatial attention. **a**, Neuronal activity into (magenta) and away from the RF (green) in the non-urgent RDM task (*n* = 51). Same data as in Fig. 2c, but restricted to the period following cue onset. **b**, Spatial signal magnitude, *S*_*ROC*_, as a function of time, for the data in **c**. Arrow marks approximate onset of differentiation. Same data as in Fig. 2d, but over a restricted time period. **c, d**, As in **a, b**, but for the non-urgent color discrimination task (*n* = 43). Same data as in Fig. 4c, d, but over a restricted time period. Note the change in activity in the 50–150 ms following cue onset (gray, shaded areas): after the motion stimulus appears near fixation (panel **a**), the LIP activity associated with the choice-target locations decreases slightly, as found in previous studies^4,5,8,9,11^; in contrast, after the color change of the choice targets (panel **c**), activity clearly increases. The decrease in **a** is maximal ∼90 ms after cue onset, at the same time that (uninformed) saccades to the choice targets are completely suppressed in the urgent CRDM task (Fig. S4). Interpreting the LIP activity as an attention signal, the motion stimulus near fixation acts as if to suppress attention at the choice targets, whereas the color change at those targets acts as if to increase the intensity of the existing signal.

**Figure S7.**
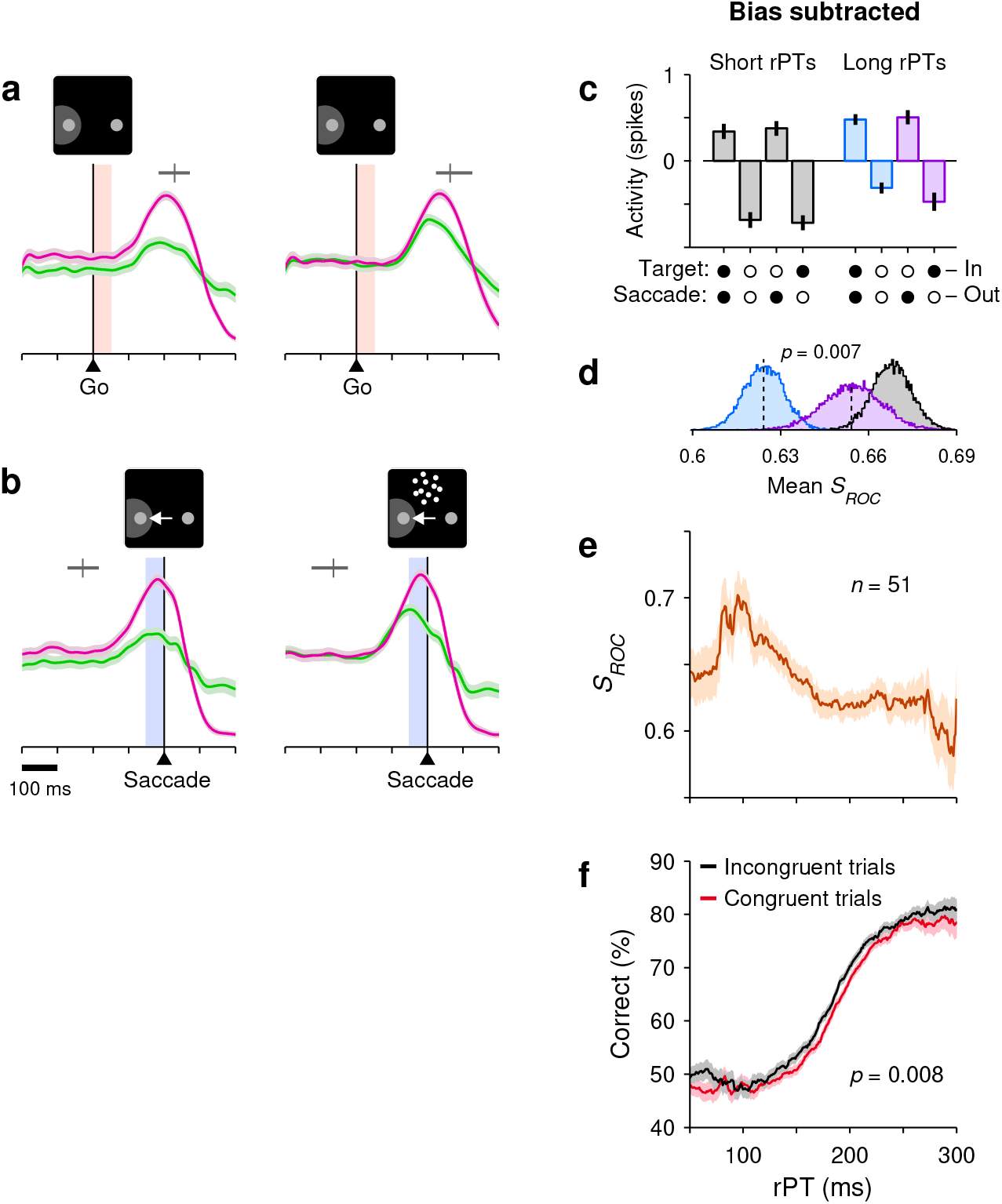
An early choice bias does not account for the results in the CRDM task. **a**, Mean, normalized LIP activity recorded in the CRDM task (*n* = 51) aligned on the go signal. Colors indicate saccadic choices into (pink) or away from the RF (green), with trials sorted into guesses (left, rPT ≤ 150 ms) and fully informed discriminations (right, rPT ≥ 200 ms). The horizontal gray bar marks the saccade onset (90% range and median). To quantify the early bias in each trial, spikes were counted in the 50 ms time window immediately following the go signal (shaded areas). **b**, Mean, normalized LIP activity recorded in the CRDM task aligned on saccade onset. Same data as in Fig. 2e, f; same trials as in **a**, but aligned differently. The horizontal gray bar marks the go signal (90% range and median). To quantify the presaccadic activity in each trial, spikes were counted in the 50 ms time window immediately preceding the saccade onset (shaded areas). **c**–**f**, Analysis results obtained after subtracting the early bias. The neural response in each trial was equal to the spike count from the standard presaccadic window (blue shade in **b**) minus the spike count from the earlier bias window (red shade in **a**). **c**, Mean normalized responses sorted by outcome, as in Fig. 3a. **d**, Mean differential signal magnitudes for the three conditions in panel **c** indicated by color, as in Fig. 3b. **e**, Neuronal performance curve showing *S*_*ROC*_ (mean ± 1 SE across trials) as a function of rPT, as in Fig. 2h. **f**, Performance in the CRDM task conditioned on neuronal activity, as in Fig. 3c. The spatial signal based on the bias-subtracted spike counts demonstrated the all same trends found originally with the unaltered presaccadic responses.

**Figure S8.**
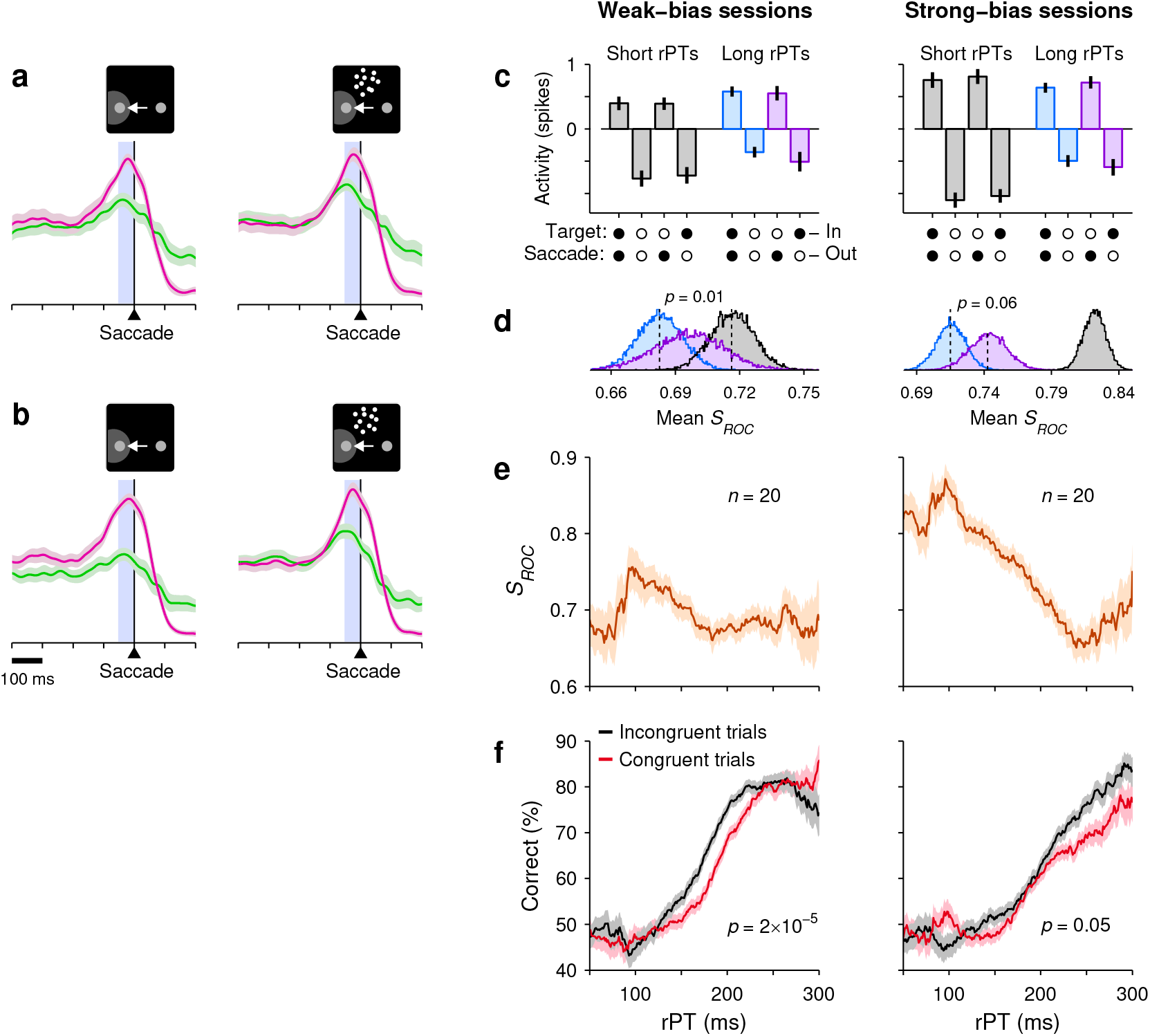
The correlation between LIP activity and CRDM behavior is qualitatively the same for sessions in which a weak or a strong choice bias was observed. For each experimental session, the early bias was quantified by counting the spikes evoked in the 50 ms immediately following the go signal (Fig. S7a, shaded areas), and calculating a spatial discrimination index (*S*_*ROC*_) using all the spike counts from that session. Based on this index, the 20 sessions with the weakest bias and the 20 with the strongest were identified, and analyses were run separately for the two groups. **a**, Mean, normalized LIP activity as a function of time in the sessions with the weakest early bias. Colors indicate saccadic choices into (pink) or away from the RF (green), with trials sorted into guesses (left, rPT ≤ 150 ms) and fully informed discriminations (right, rPT ≥ 200 ms). **b**, As in **a**, but for the sessions with the strongest early bias. **c**–**f**, Analysis results for weak- (left column) and strong-bias sessions (right column). **c**, Mean normalized responses sorted by outcome, as in Fig. 3a. **d**, Mean differential signal magnitudes for the three conditions in panel **c** indicated by color, as in Fig. 3b. **e**, Neuronal performance curves showing the presaccadic *S*_*ROC*_ (mean ± 1 SE across trials) as a function of rPT, as in Fig. 2h. **f**, Performance conditioned on neuronal activity, as in Fig. 3c. Although the magnitude of the spatial signal before the saccade did vary with the magnitude of the early bias, stronger presaccadic differentiation was still associated with shorter processing times and poorer performance, regardless of the bias.

**Figure S9.**
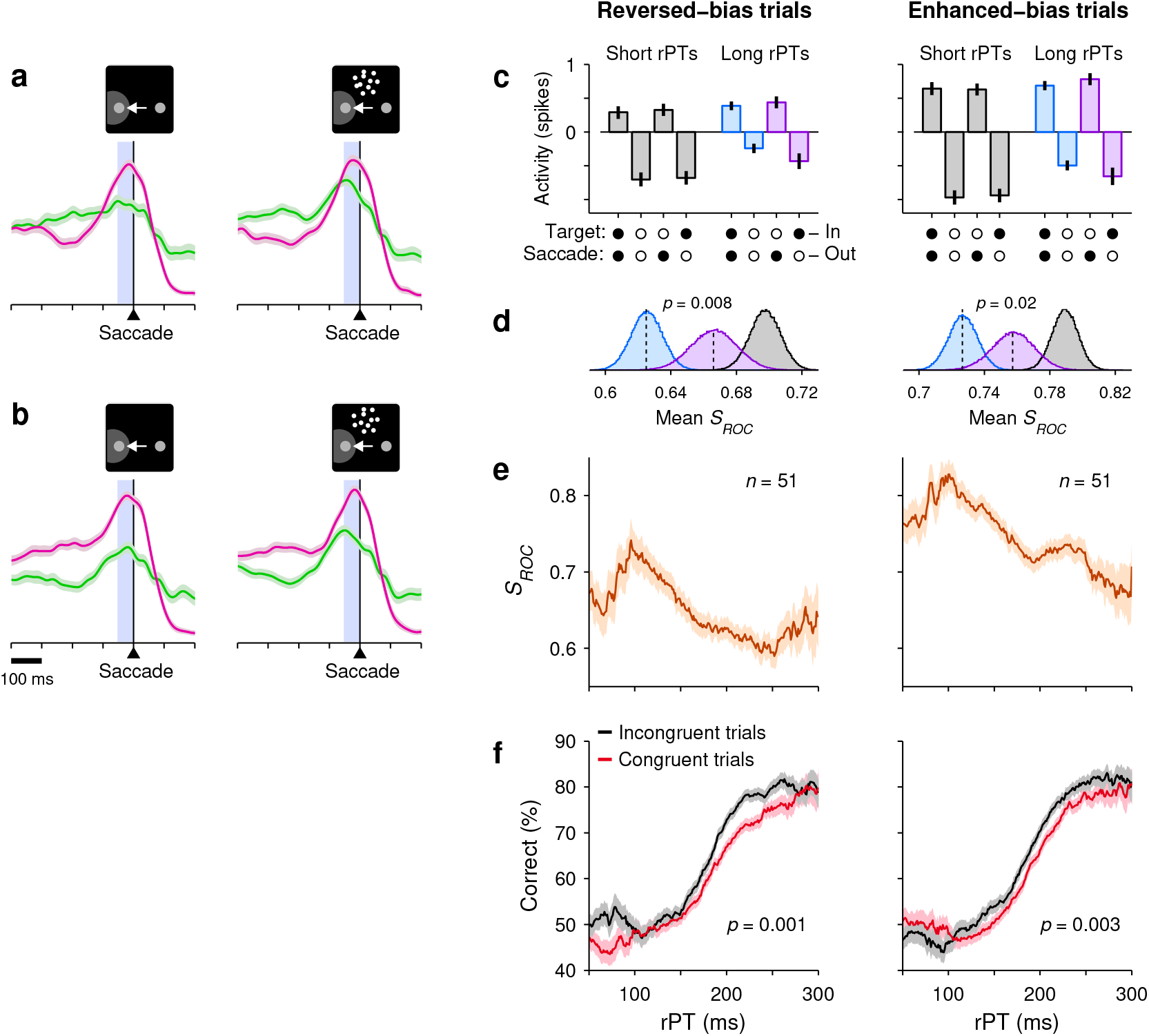
Creating artifical biases through trial sorting does not change the observed correlation between LIP activity and CRDM behavior. For each recorded neuron, the early bias in each trial was quantified by counting the spikes evoked in the 50 ms immediately following the go signal (Fig. S7a, shaded areas). Then, trials into and away from the RF were separately split into two groups according to the median spike count in the bias window, and the data for each of the 4 resulting groups were pooled across neurons. Finally, two such groups of trials (strong response in; weak response away) were paired to create a data set with an enhanced bias into the RF, and the other two groups (weak response in; strong response away) were paired to create a data set with a reversed bias, i.e., a bias away. Analyses were then run on the two halfs of the data thus parsed. **a**, Mean, normalized LIP activity as a function of time for the data set with a reversed bias initially favoring the away direction. Colors indicate saccadic choices into (pink) or away from the RF (green), with trials sorted into guesses (left, rPT ≤ 150 ms) and fully informed discriminations (right, rPT ≥ 200 ms). **b**, As in **a**, but for the data set with an enhanced early bias toward the RF. **c**–**f**, Analysis results for reversed-(left column) and enhanced-bias data sets (right column). Same format as in Fig. S8c–f. Sorting trials in this fashion strongly alters the starting point of the evoked presaccadic responses, but the subsequent changes in activity maintain a consistent qualitative relationship with processing time and choice outcome, regardless of that initial condition.

**Figure S10.**
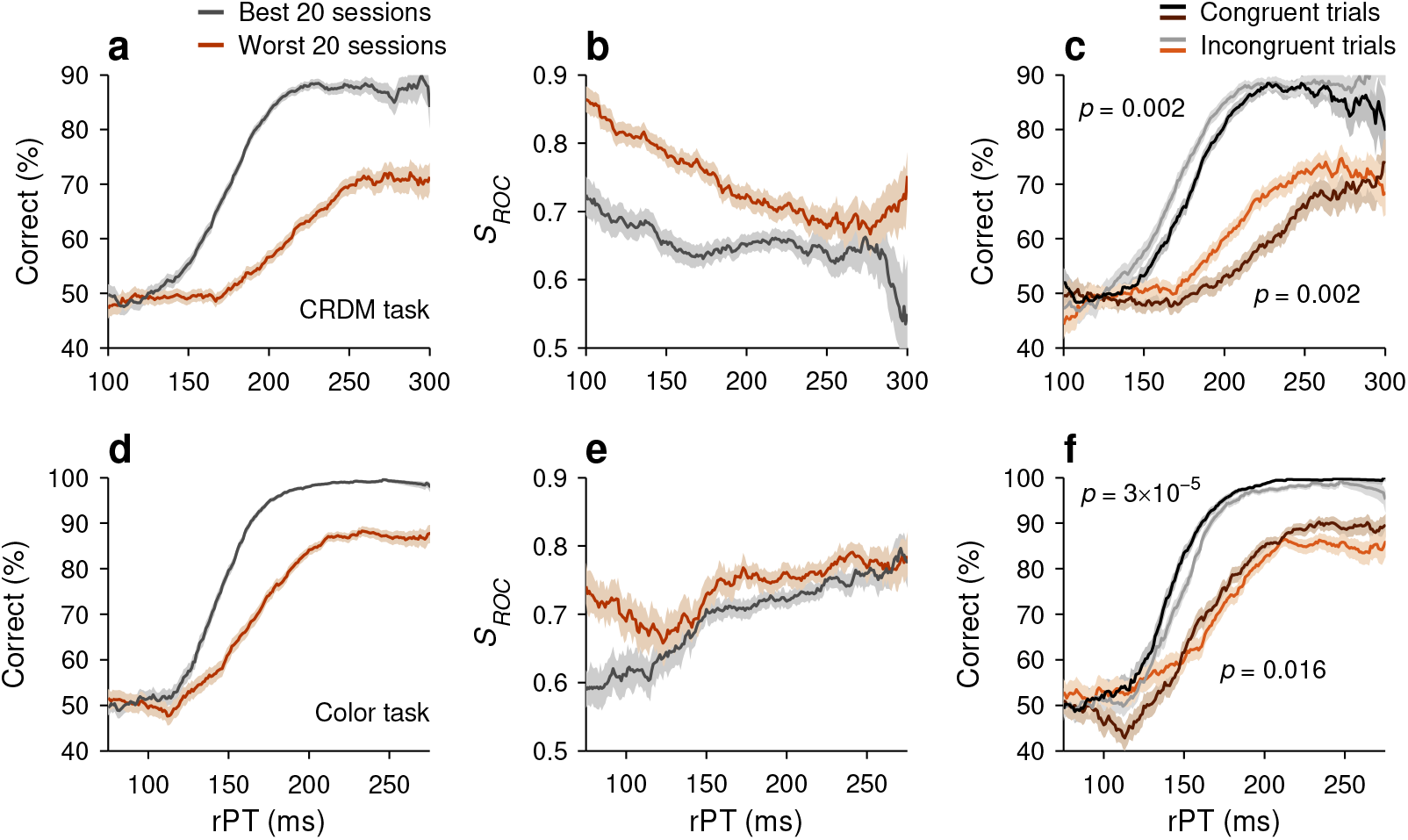
Modulation of LIP activity during high-versus low-performance sessions. For both the motion- and color-based urgent tasks, recording sessions were ranked according to overall percent correct. The 20 sessions with the best performance (gray traces) and the 20 sessions with the poorest performance (brown traces) were selected, and analyses were run for each group separately. **a**, Tachometric curves, i.e., percent correct as a function of processing time, for the best and worst CRDM sessions. **b**, Neurometric curves in the CRDM task, i.e., magnitude of presaccadic differentiation as a function of processing time (as in Fig. 2h). In both groups of sessions, stronger LIP differentiation was associated with less evidence (shorter rPTs). **c**, Tachometric curves conditioned on neuronal activity in the CRDM task (as in Fig. 3c). In both groups of sessions, stronger LIP differentiation (congruent condition) was associated with worse perceptual performance. **d**– **f**, Same as **a**–**c**, but for the urgent color discrimination task (as in Figs. 4h and 3f). In this task, stronger LIP differentiation was associated with more evidence (longer rPTs; **e**) and improved perceptual performance (**f**). In each task, the relationship between neuronal activity in LIP and behavior was similar for the two groups of sessions, in spite of the dramatically different performance levels.

**Figure S11.**
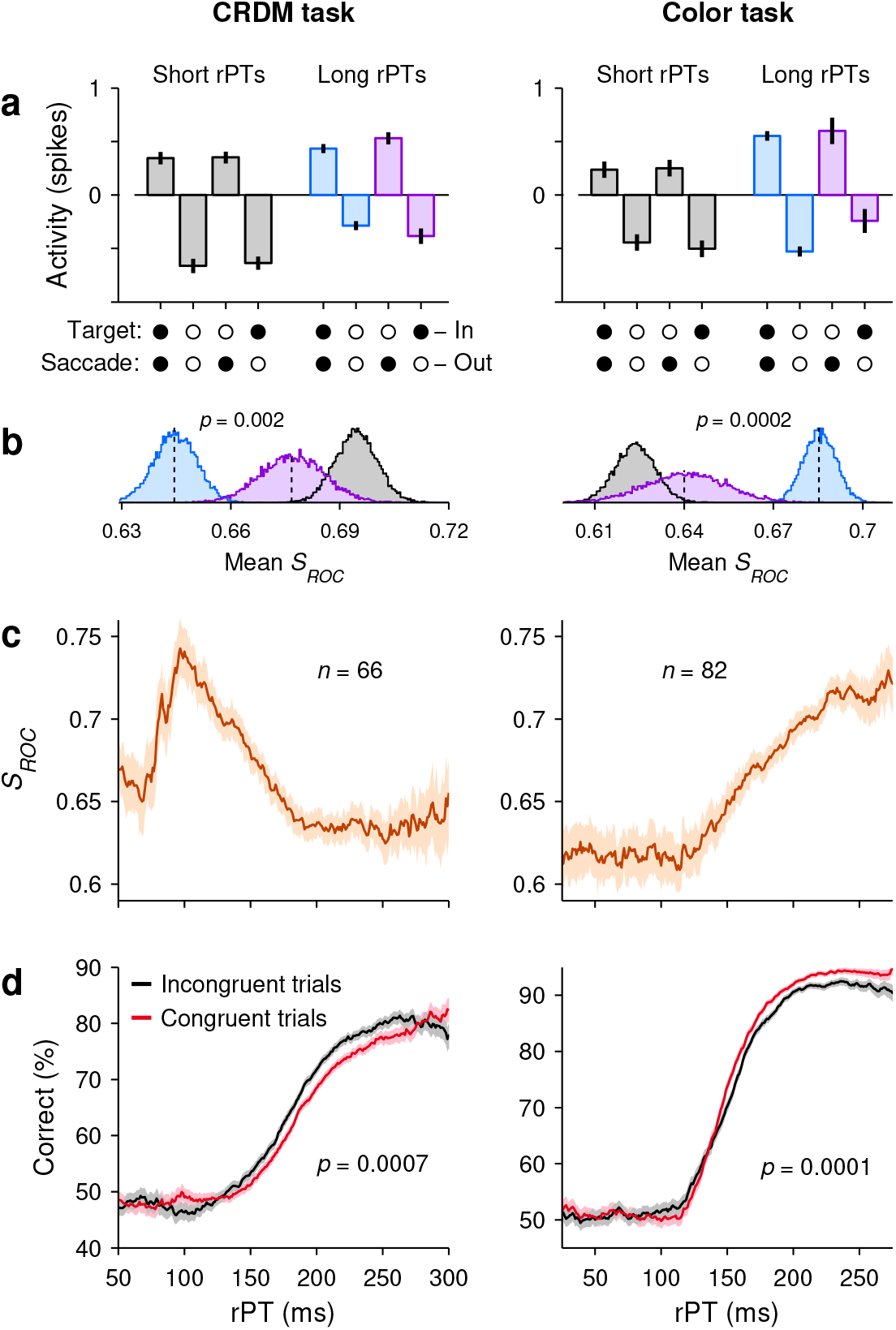
Results with extended neuronal populations. The results in the main figures were based on neurons (*n* = 51 for the CRDM task, *n* = 56 for the urgent color discrimination task) that were fully characterized and satisfied several criteria: they had both visually-driven and saccade-related responses, and their RFs were well defined and consistent across tasks (Methods). This resulted in the exclusion of 15 additional neurons recorded in the CRDM experiment and 26 in the color-based. Here we show the results of analyses that included all the neurons recorded in the CRDM (left column, *n* = 66) and the urgent color discrimination task (right column, *n* = 82), with no exclusions. **a**, Mean, normalized neuronal activity pooled across neurons and sorted by experimental condition (as in Fig. 3a, d). **b**, Distributions for mean presaccadic differentiation values (*S*_*ROC*_), obtained by bootstrapping, for each of the three conditions above, as indicated by the corresponding colors (as in Fig. 3b, e). **c**, Neurometric curves, i.e., magnitude of presaccadic differentiation as a function of processing time (as in Figs. 2h, 4h). **d**, Behavioral performance conditioned on neuronal activity (as in Fig. 3c, e). Inclusion of the additional populations did not alter the results substantially.

**Figure S12.**
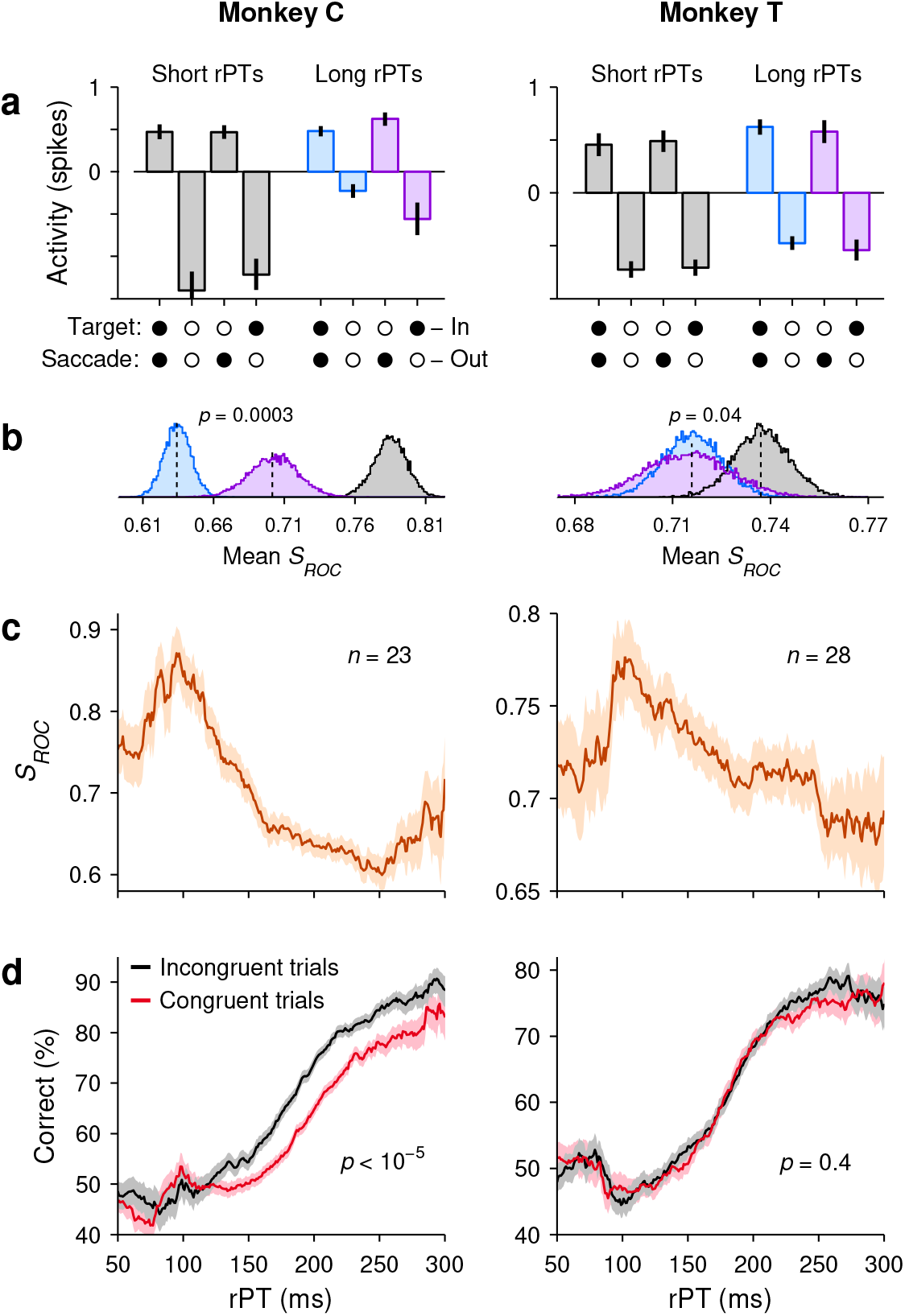
Key results in the CRDM task computed separately for monkeys C (left column, *n* = 23) and T (right column, *n* = 28). **a**, Normalized LIP activity during guesses (rPT ≤ 150 ms, gray) and informed choices (rPT *>* 150 ms, blue, purple) sorted by experimental condition (as in Fig. 3a). **b**, Distributions for mean presaccadic differentiation values (*S*_*ROC*_) for each of the three conditions above, as indicated by color (as in Fig. 3b). **c**, Neurometric curve, i.e., magnitude of pre-saccadic differentiation as a function of processing time (as in Fig. 2h). **d**, Behavioral performance conditioned on neuronal activity (as in Fig. 3c).

**Figure S13.**
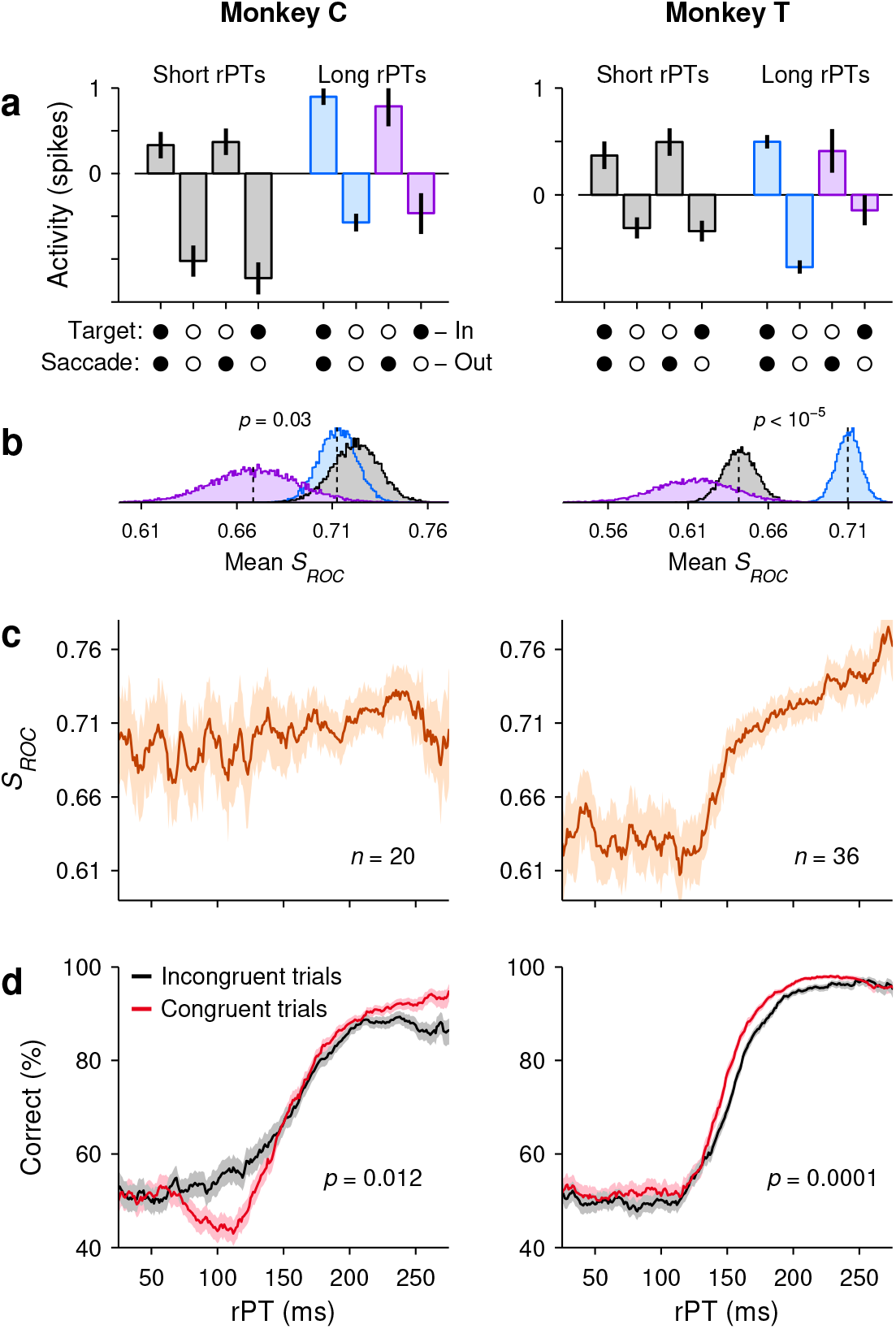
Key results in the urgent color discrimination task computed separately for monkeys C (left column, *n* = 20) and T (right column, *n* = 36). **a**, Normalized LIP activity during guesses (rPT ≤ 125 ms, gray) and informed choices (rPT *>* 125 ms, blue, purple) sorted by experimental condition (as in Fig. 3d). **b**, Distributions for mean presaccadic differentiation values (*S*_*ROC*_) for each of the three conditions above, as indicated by color (as in Fig. 3e). **c**, Neurometric curve, i.e., magnitude of presaccadic differentiation as a function of processing time (as in Fig. 4h). **d**, Behavioral performance conditioned on neuronal activity (as in Fig. 3f).

**Figure S14.**
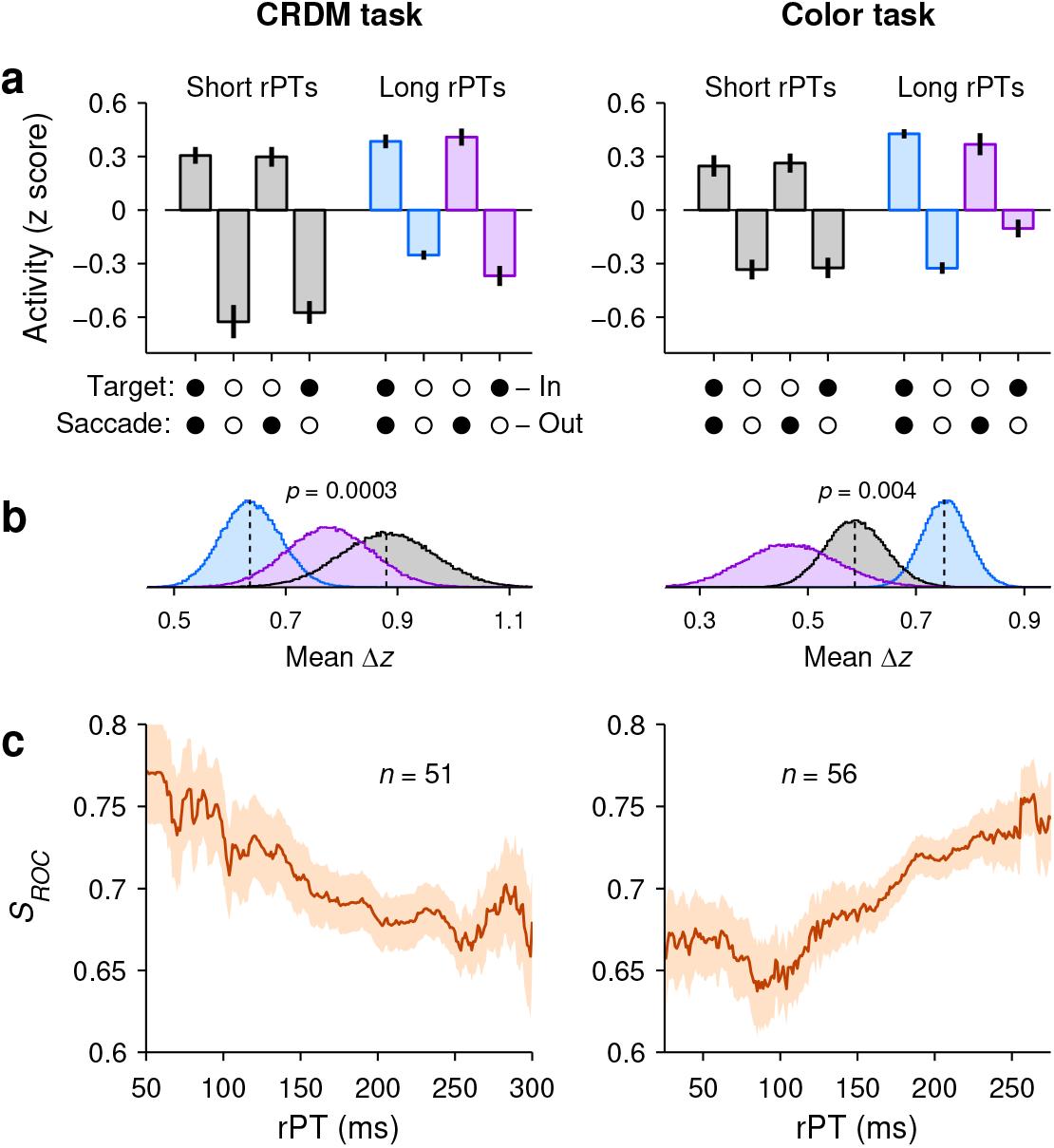
Alternate procedure for combining the data across neurons. In the main figures (Figs. 2h, 4h, 3a, d), we first pooled all the spike counts from all the neurons, separately for saccades into (*S*_*in*_) and away from the RF (*S*_*out*_), and then quantified the separation between the two resulting distributions by computing the *S*_*ROC*_ (Methods; Fig. S15). Here we present an alternate analysis in which *S*_*in*_ and *S*_*out*_ conditions are first contrasted for each neuron and then the results are averaged across neurons. Results are for the CRDM (left column, *n* = 51) and the urgent color discrimination task (right column, *n* = 56). **a**, LIP activity during guesses and informed choices sorted by outcome (same short- and long-rPT intervals as in Fig. 3a, d). Activity corresponds to presaccadic spike counts (same as in other analyses) that were z-scored for each neuron and then averaged across neurons. Data are mean ± 1 SE across cells. **b**, Differential signal magnitudes for the three conditions in **a** indicated by color. Here, the differential signal of each cell is the mean difference between the z-scored responses in *S*_*in*_ and *S*_*out*_ conditions. Thus, the mean Δ*z* is equal to ⟨*z*_*IN*_ − *z*_*OUT*_ ⟩, where the brackets indicate an average over neurons. Curves are bootstrapped distributions. **c**, Mean neurometric curve, i.e., magnitude of presaccadic differentiation as a function of processing time. In this case, a neurometric curve was first computed for each individual neuron (bin width = 81 ms) and then the results were averaged over neurons. Shades indicate ± 1 SE across cells. The results of these analyses are more variable than those in the main figures, but show the same qualitative trends for how spatial discriminability depends on processing time and trial outcome in each task.

**Figure S15.**
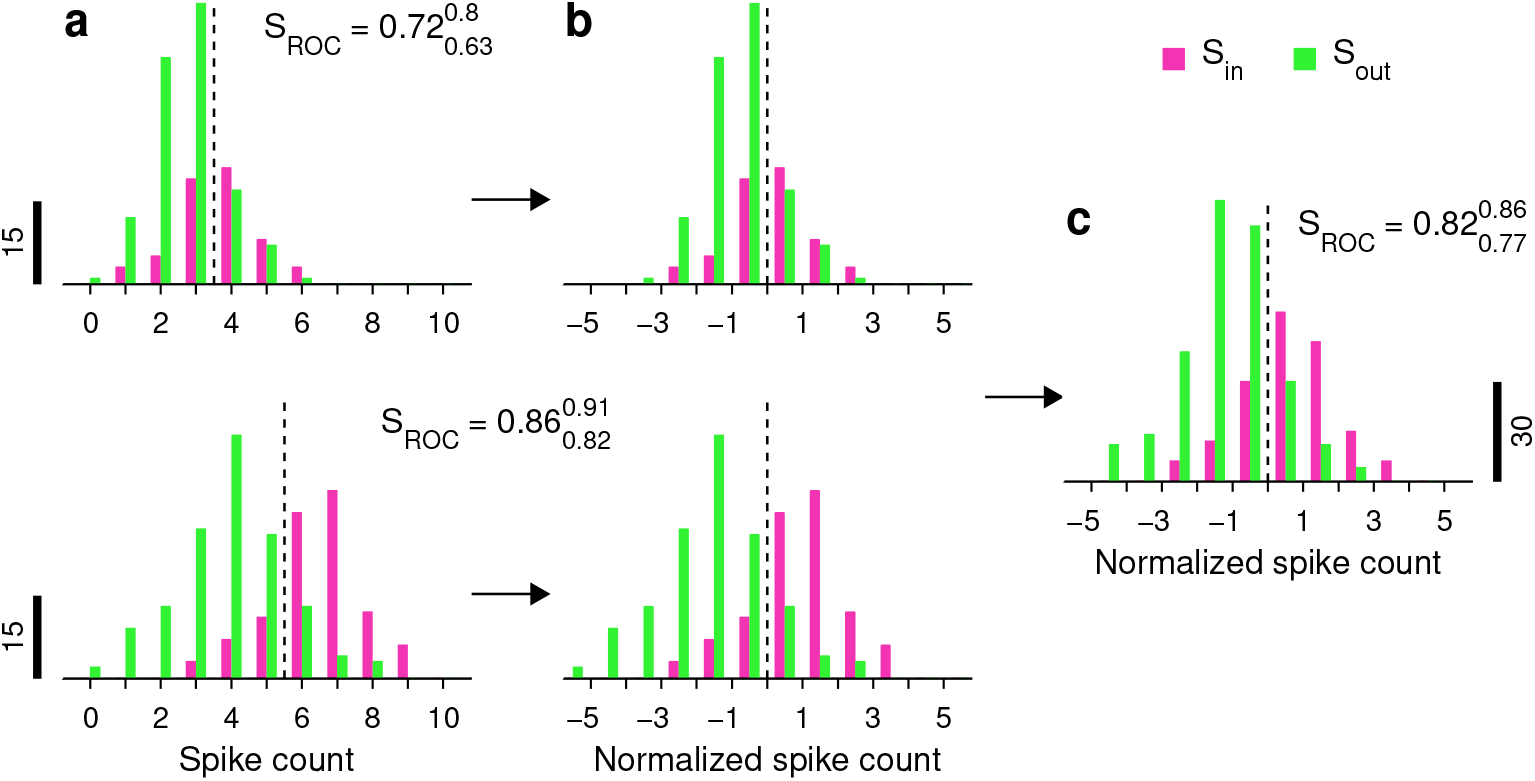
Procedure for pooling the data across neurons before computing the average magnitude of their spatial signal. This example illustrates the pooling method for two neurons recorded in the CRDM task. For each cell, the response in each trial was the spike count collected in the 50 ms immediately preceding saccade onset. Responses were sorted by condition, for trials in which the saccade was into the RF (*S*_*in*_, pink bars) and for trials in which the saccade was away (*S*_*out*_, green bars). In this case all rPTs are included. **a**, Spike count histograms from two LIP neurons. For each cell, the dashed line is the value (*θ*) intermediate between the mean spike counts for *S*_*in*_ and *S*_*out*_ trials. The magnitude of the differential response based on each cell’s data is indicated, with 95% confidence limits (from bootstrap). **b**, Same data as in **a**, but after having subtracted *θ* from each spike count. Individual *S*_*ROC*_ values do not change, as they are invariant to linear transformations of the data. **c**, Histograms for the pooled, normalized data from the two neurons. Note that the resulting *S*_*ROC*_, which is computed exactly as for the single cells, is intermediate between their values.

